# GABA signalling in guard cells acts as a ‘stress memory’ to optimise plant water loss

**DOI:** 10.1101/2019.12.22.885160

**Authors:** Bo Xu, Yu Long, Xueying Feng, Xujun Zhu, Na Sai, Larissa Chirkova, Johannes Herrmann, Mamoru Okamoto, Rainer Hedrich, Matthew Gilliham

## Abstract

The non-protein amino acid γ-aminobutyric acid (GABA) has been proposed to be an ancient messenger for cellular communication conserved across biological kingdoms. GABA has well-defined signalling roles in animals; however, whilst GABA accumulates in plants under stress it has not been determined if, how, where and when GABA acts as an endogenous plant signalling molecule. Here, we establish that endogenous GABA is a *bona fide* plant signal, acting via a mechanism not found in animals. GABA antagonises stomatal movement in response to opening and closing stimuli in multiple plant families including dicot and monocot crops. Using *Arabidopsis thaliana*, we show guard cell GABA production is necessary and sufficient to influence stomatal aperture, transpirational water loss and drought tolerance via inhibition of stomatal guard cell plasma membrane and tonoplast-localised anion transporters. This study proposes a novel role for GABA – as a ‘stress memory’ – opening new avenues for improving plant stress tolerance.

---

The regulation of stomatal pore aperture is a key determinant of plant productivity and drought tolerance, and profoundly impacts climate due to its influence on global carbon and water cycling^1, 2, 3^. The stomatal pore is delineated by a guard cell pair. Fine control of ion and water movement across guard cell membranes, via transport proteins, determines cell volume and pore aperture following opening and closing signals such as light and dark^2, 4, 5^ (Fig. 1a, b). Due to their critical roles, and their ability to respond to and integrate multiple stimuli, guard cells have become a preeminent model system for investigating plant cell signalling^6^ resulting in the elucidation of many critical pathways involved in plant biotic and abiotic stress tolerance^7, 8, 9^.

**Figure 1.**
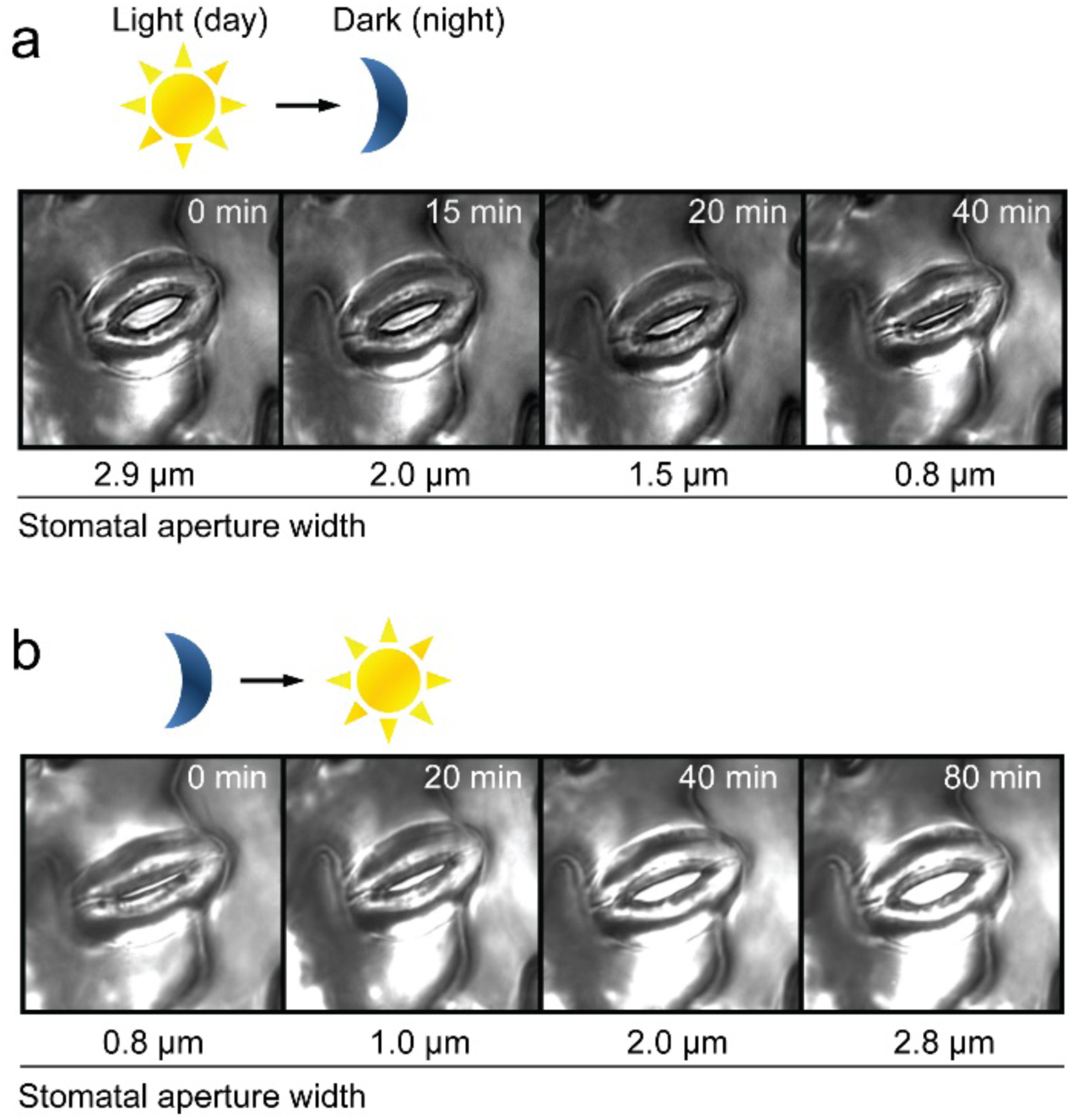
Guard cells respond to light signals. **a-b**,Time-course of dark-induced stomatal closure (**a**) and light-induced stomatal opening (**b**) with actual stomatal aperture width indicated below; light-to-dark transition mimics day-to-night transition which closes stomata (**a**) and dark-to-light transition mimics night-to-day transition which opens stomata (**b**), light intensity,150 µmol m^-2^ s^-1^.

GABA signalling in mammals relies upon receptor-mediated polarization of neuronal cell membranes^10, 11^. Speculation that GABA could be a signal in plants is decades old^12^, but a definitive demonstration of its mode of action remains elusive. The discovery that the activity of Aluminium-activated Malate Transporters (ALMTs) can be regulated by GABA^13^ represents a putative mechanism by which GABA signals could be transduced in plants. Stomatal guard cells contain a number of ALMTs that impact stomatal movement and transpirational water loss^14, 15, 16^. Therefore, guard cells represent an ideal system to test whether GABA signalling occurs in plants.

In this study, we give the first definitive demonstration that GABA is legitimate plant signalling molecule by showing the mechanism by which it acts in stomatal guard cells. Although guard cell signalling is relatively well defined^6, 17^, our work uncovers a new pathway regulating plant water loss. Significantly, we show that GABA does not initiate stomatal movement, rather it antagonises movement via negative regulation of guard cell plasma membrane and tonoplast ion transport. Specifically, we find that GABA increases under a water deficit and inhibits ALMT9 activity. ALMT9 is a major pathway for mediating anion uptake into the vacuole during stomatal opening; its inhibition by GABA reduces stomatal opening during a water deficit which reduces transpirational water loss and confers improved plant drought tolerance. As such, we propose that GABA is a prime candidate molecule to confer a ‘memory’ of water stress in guard cells^18^.

## Results

### GABA antagonises stomatal movements

To validate whether GABA is a physiological signal that modulates stomatal movement, our initial experiments used excised *Arabidopsis thaliana* epidermal strips where guard cells are directly accessible to a chemical stimuli^8, 19, 20, 21^. Interestingly, when applied under constant light or dark conditions, exogenous GABA or its analog muscimol^22^ did not elicit a change in stomatal aperture. Instead we found that both compounds suppressed dark-induced stomatal closure and light-induced opening (Fig. 2a, b; Supplementary Fig. 1a-b). We observed the same behaviour in intact leaves. In response to a dark-to-light transition, leaves fed muscimol through the petiole dampened gas exchange rates, consistent with a slower rate of stomatal opening (Supplementary Fig. 1c-e). We tested whether our results could be explained by GABA or muscimol treatment killing guard cells, which prevented their movement, by performing a washout experiment on epidermal strips. As would be expected from viable cells, after removal of the GABA or muscimol treatment we found that stomatal guard cells responded to a dark treatment by closing the stomatal pore (Supplementary Fig. 2a, b); controls that received constant light remained open (Supplementary Fig. 2a, b). Collectively, these data indicate that GABA signals would likely act to modulate stomatal movement in the face of a stimulus rather than stimulating a transition itself.

**Figure 2.**
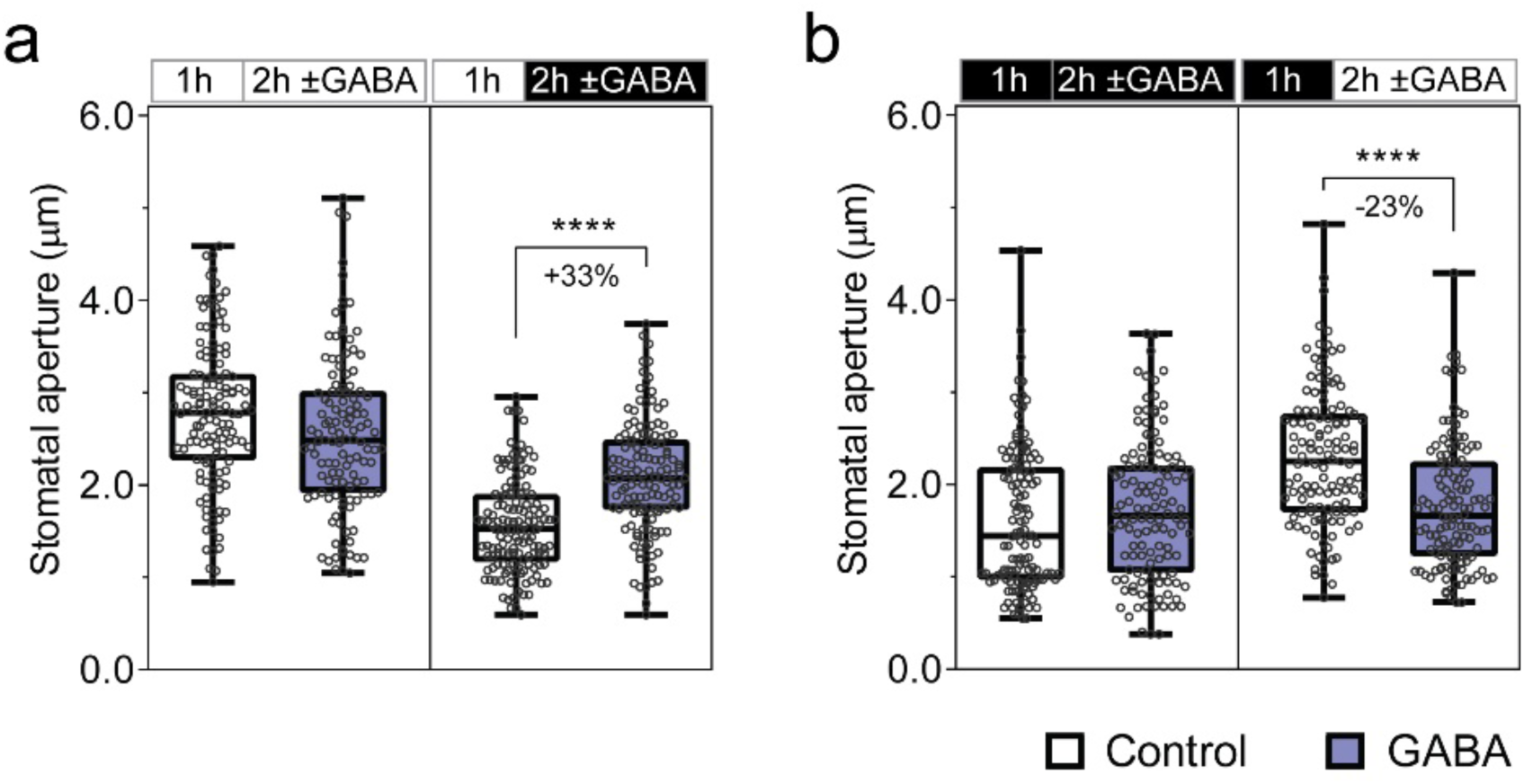
Exogenous GABA application antagonises stomatal movement. **a-b**, Stomatal aperture measurement of wildtype *A. thaliana* leaves in response to light or dark. Epidermal strips were pre-incubated in stomatal measurement buffer for 1 h under light (**a**) or dark (**b**), followed by 2 h incubation under constant light (**a**), dark (**b**), light-to-dark transition (**a**) or dark-to-light transition (**b**) as indicated above graphs by black (dark) or white (light) bars, together with the application of 2 mM GABA; n = 129 for control (constant light), n = 121 for GABA (constant light), n = 137 for control (light-to-dark transition) and n = 135 for GABA (light-to-dark transition) (**a**); n = 122 for control (constant dark), n = 124 for GABA (constant dark), n = 123 for control (dark-to-light transition) and n = 130 for GABA (dark-to-light transition) (**b**); all experiments were repeated at least twice from different batches of plants with blind treatments. All data are plotted (box plots contain five horizontal lines representing - from top to bottom - maximum, second quartile, median, third quartile and minimum values); statistical difference was determined by Student’s *t*-test, \*\*\*\**P*< 0.0001.

### GABA is a universal stomatal behaviour modifier in valuable crops

To test whether GABA is a universal modulator of stomatal control, we explored whether GABA or muscimol treatment of epidermal strips attenuated stomatal responses of other plant species to light or dark transitions, including the dicot crops *Vicia faba* (broad bean), *Glycine max* (soybean) and *Nicotiana benthamiana* (tobacco-relative), and the monocot *Hordeum vulgare* (barley) (Supplementary Fig. 3). The widespread inhibition of movement observed suggests that GABA is a universal ‘brake’ for stomatal movement in plants.

### GABA accumulation in guard cells contributes to the regulation of transpiration and drought performance

Stomatal control is explicitly linked with the regulation of plant water loss, which impacts the survival of plants under drought^7^; the wider the stomatal aperture, the greater the water loss of plants, the poorer the survival of plants under a limited water supply. As GABA has often been found to accumulate under stress^23^ and modulates the extent of opening and closure (Fig. 2), we examined the hypothesis that endogenous GABA might increase in concentration under a water deficit and act as a signal. In wildtype plants, a drought treatment was applied by withholding watering. This resulted in the gradual depletion of soil gravimetric water and a reduction in leaf relative water content (RWC) (Supplementary Fig. 4a, b). GABA accumulation in drought stressed leaves increased by 35% compared to that of well-watered leaves (water *vs*. drought at 7 d: 1.07 ± 0.08 vs. 1.44 ± 0.11 nmol mg^-1^ FW) (Supplementary Fig. 4c).

Arabidopsis T-DNA insertional mutants for the major leaf GABA synthesis gene, *Glutamate Decarboxylase 2* (*GAD2)* (*gad2-1* and *gad2-2*) had >75% less GABA accumulation in leaves than in wildtype plants, whilst concentrations in roots were unchanged (Fig. 3a; Supplementary Fig. 4d-f). Furthermore, leaves of *gad2* plants did not accumulate GABA under drought conditions unlike wildtype controls where GABA increased by 45% after 3 days, which was maintained at this elevated level after 7 days drought (Fig. 3a). Under standard conditions both *gad2* mutant lines exhibited greater stomatal conductance than wildtype plants due to wider stomatal pores, with stomatal density being identical to wildtype levels (Fig. 3b; Supplementary Fig. 4g, h). The application of exogenous GABA to *gad2* leaves reduced the extent of their stomatal opening to that of wildtype plants without GABA treatment (Supplementary Fig. 4i, j). This indicated that *gad2* stomata would be competent in a GABA response if sufficient GABA was present.

**Figure 3.**
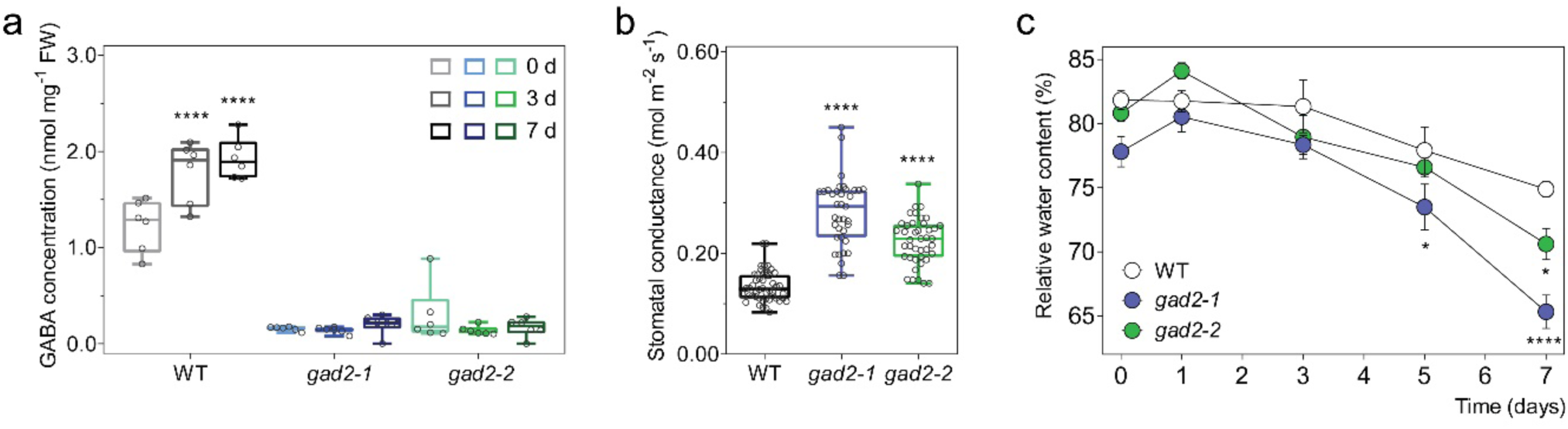
Leaf GABA regulates transpiration. **a**, Leaf GABA concentration of 5-week old wildtype (WT), *gad2-1* and *gad2-2* plants following drought treatment for 0 d, 3 d and 7 d, n = 6. **b**, Stomatal conductance of Arabidopsis WT (n = 48), *gad2-1* (n = 37) and *gad2-2* (n = 41) plants determined by AP4 Porometer, data collected from three independent batches of plants. **c**, Relative leaf water content of WT, *gad2-1* and *gad2-2* plants following drought treatment for 0 d, 1 d, 3 d, 5 d, and 7 d, n = 6. All data are plotted (**a, b**) or represented as mean ± s.e.m (**c**); statistical difference was determined using Two-way ANOVA (**a, c**) or One-way ANOVA (**b**); \**P*< 0.05 and \*\*\*\**P*< 0.0001.

Under drought the leaf RWC of *gad2* plants lowered more quickly than in wildtype (Fig. 3c). However, transcriptional profiles of key ABA-marker and GABA-related genes were similar in wildtype and *gad2* lines (except for *GAD2*), indicating no detectable differential induction of these pathways (Supplementary Fig. 5). These results confirm that *GAD2* is critical for leaf GABA production under stress, and for regulating plant water loss and drought tolerance^24^. Furthermore, *GAD2* transcription and GABA accumulation exhibit diurnal regulation, except during stress when both are constitutively high^25^. GABA usually peaks at the end of the dark cycle prior to stomatal opening and reaches a minimum when stomatal conductance is at its maximum near subjective midday^25^. These data are consistent with GABA not only playing a role to slow and minimise stomatal opening under stress, but also indicates that GABA could fulfil this role under non-stressed conditions. The high stomatal conductance phenotype of the *gad2* plants under well-watered conditions would also seem to indicate that GABA is a vital component in regulating stomatal control and water loss even under non-watered limited conditions.

The expression pattern of *GAD2* was examined. Histochemical staining corroborated that *GAD2* is highly expressed in leaves, particularly in guard cells (Supplementary Fig. 6a, b). GAD2 is a cytosolic enzyme^26^. To examine if cytosolic GABA biosynthesis within the guard cell was sufficient to modulate transpiration we expressed - specifically in the guard cell^27^ - a constitutively active form of *GAD2* (*GAD2Δ*) ^26, 28^. This led to a large increase in leaf GABA accumulation (Fig 4a), and to complementation of the steady-state stomatal conductance and aperture phenotypes of *gad2* plants to wildtype levels (Fig 4b, c; Supplementary Fig. 6c, d); no change in stomatal density was detected (Supplementary Fig. 6e). The faster opening and closure kinetics of *gad2-1* were recovered to wildtype-like rates by guard-cell specific expression of *GAD2Δ* (Fig. 4d, e). Drought-sensitivity of *gad2* plants, compared to wildtype, was also abolished by guard cell specific expression of *GAD2Δ* (Fig. 4f, g). In contrast, when *GAD2Δ* was specifically expressed in *gad2* spongy mesophyll^29^, we detected a significant increase in leaf GABA but no change in stomatal conductance (Supplementary Fig. 7a-d). In a further experiment we expressed full length *GAD2* under a guard cell-specific promoter (*gad2/GC1∷GAD2*). This did not complement the high stomatal conductance of the *gad2-1* line to wildtype levels under standard conditions, whereas its constitutive expression (driven by pro35S-CAMV) did (Supplementary Fig. 7e-j). Interestingly, under drought *gad2-1/GC1∷GAD2* lines reduced their stomatal conductance significantly more than that of *gad2-1* plants and had similar leaf RWC comparable to wildtype plants following 5 days of drought (Supplementary Fig. 7e-h). This suggests that the GAD2 regulatory domain removed in GAD2Δ is important in stimulating GABA production under drought. Collectively, these data demonstrate that guard-cell specific cytosolic GABA accumulation is sufficient for controlling stomatal aperture and transpiration under drought, but suggests a role for other cell-types in fine tuning GABA signals under standard conditions.

**Figure 4.**
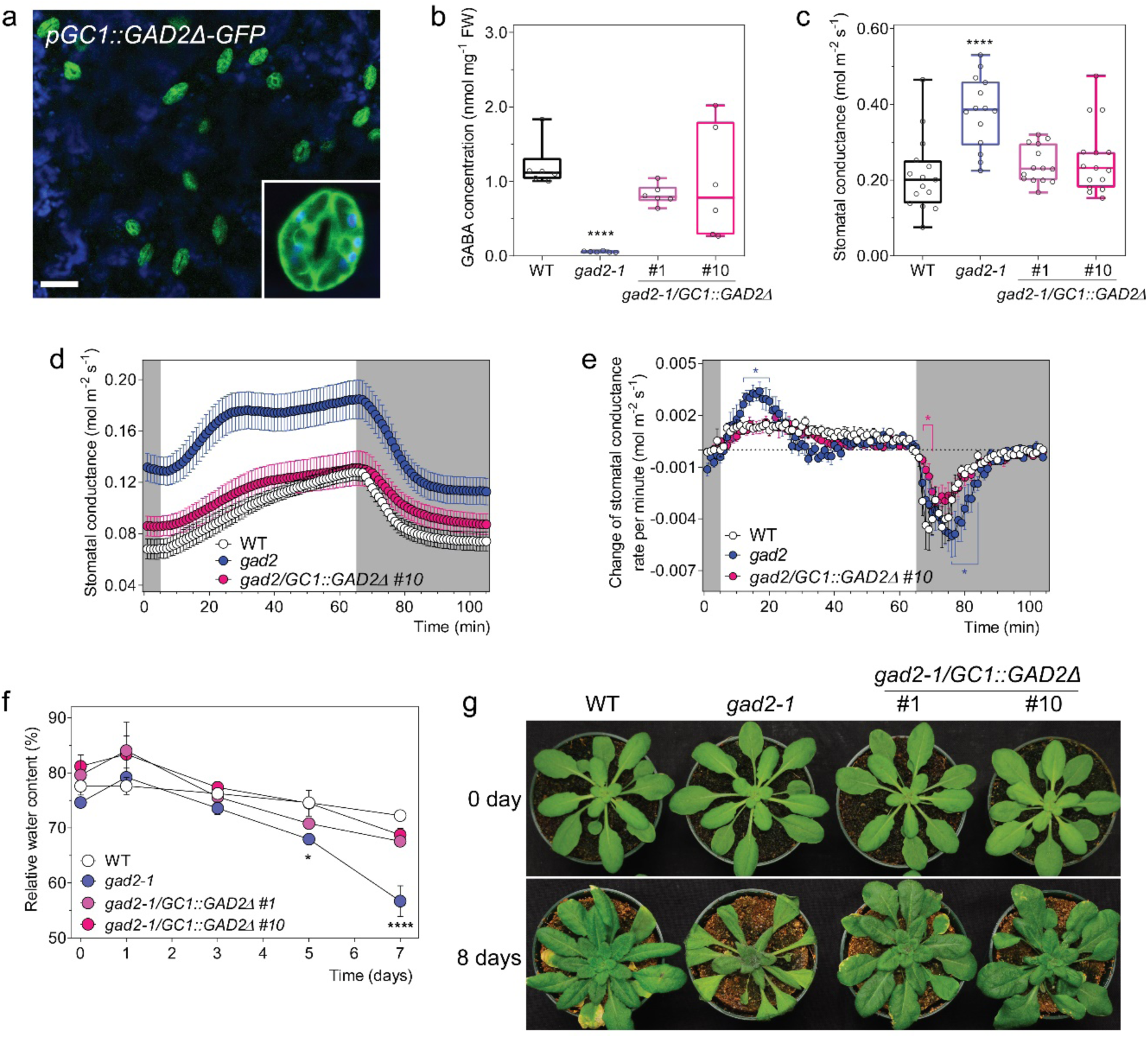
Guard-cell GABA regulates water loss and drought tolerance. **a**, Confocal images of *gad2*-1 plants expressing *GC1∷GAD2Δ*-*GFP* (*gad2-1/GC1∷GAD2Δ*-*GFP*); GFP fluorescence and chlorophyll autofluorescence (blue) of the leaf abaxial side of 3-4 week-old *gad2-1/GC1∷GAD2Δ*-*GFP* plant indicates that the *GC1* promoter drives *GAD2Δ* expression specifically in guard cells, scale bars = 50 µm (**a**). **b**, Leaf GABA accumulation of 5-6 week-old WT, *gad2-1, gad2-1/GC1∷GAD2Δ* #1 and #10 plants grown under control conditions, n= 6. **c**, Stomatal conductance of WT (n = 15), *gad2-1* (n = 14), *gad2-1/GC1∷GAD2Δ* #1 (n = 14) and #10 (n = 15) plants under control conditions, data collected from two independent batches of plants.. **d**, Stomatal conductance of WT (n = 9), *gad2-1* (n = 9) and *gad2-1/GC1∷GAD2Δ* #10 (n = 8) plants in response to dark (shaded region) and 150 µmol m^-2^ s^-1^ light (white region), measured using a LiCor LI-6400XT system. **e**, Change in stomatal conductance each minute calculated using dConductance/dt (min) of the data represented in (**d**). **f**, Relative leaf water content of WT, *gad2-1, gad2-1/GC1∷GAD2Δ* #1 and #10 plants following drought treatment for 0 d, 1 d, 3 d, 5 d, and 7d; n = 4 for 0 d, 1 d, 3 d, 5 d samples and n = 5 for 7 d samples, except that n = 3 for 0 d *gad2-1* and 1 d *gad2-1/GC1∷GAD2Δ* #1. **g**, Representative images of WT, *gad2-1, gad2-1/GC1∷GAD2Δ* #1 and #10 plants (shown in **i**) before (0 day) and after (8 days) drought treatment as indicated. Pot size 2.5 inch diameter x 2.25 inch height (LiCor). All data are plotted (**b, c**) or represented as mean ± s.e.m (**d-f**); statistical difference was determined using One-way ANOVA (**b, c**) or Two-way ANOVA (**e, f**); \**P*< 0.05 and \*\*\*\**P*< 0.0001.

When *GAD2Δ* was specifically overexpressed in the guard cells of wildtype Arabidopsis plants, the stomatal conductance of transgenic plants in standard and drought conditions was lowered (Fig 5a, b). As a consequence, the plants overexpressing *GAD2Δ* in the wildtype background maintained higher leaf RWC than wildtype plants after 10-days of drought treatment (Fig 5c, d). Furthermore, a greater percentage of plants overexpressing *GAD2Δ* in the wildtype background survived following re-watering after a 12-day drought treatment (Supplementary Fig. 8). As such, we show here that GABA overproduction can lower water loss and improve drought tolerance.

**Figure 5.**
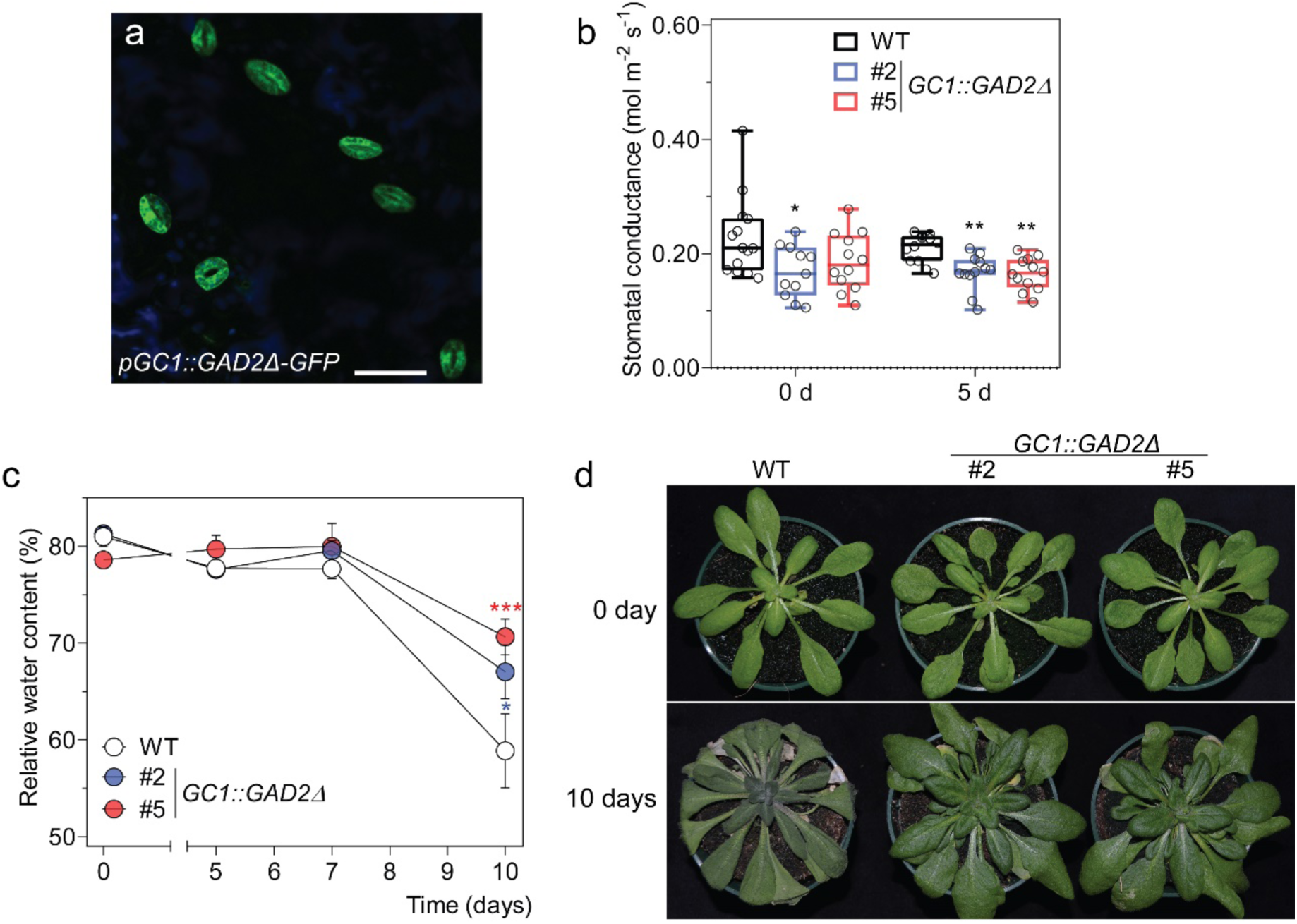
Guard-cell overexpression of *GAD2Δ* improves water-use efficiency under drought conditions. **a**, Confocal images of wildtype Arabidopsis plants expressing *GC1∷GAD2Δ*-*GFP*; GFP fluorescence and chlorophyll autofluorescence (blue) of the leaf abaxial side of 3-4 week-old plants, scale bars = 50 µm. **b**, Stomatal conductance of WT, wildtype Arabidopsis expressing *GAD2Δ* in the guard cells using *GC1* promoter – *GC1∷GAD2Δ* #2 and #5 plants before (0 d) and after (5 d) drought treatment; n = 14 for WT, n = 11 for *GC1∷GAD2Δ* #2 and n = 15 for *GC1∷GAD2Δ* #2 at 0 d; and n = 12 for WT, *GC1∷GAD2Δ* #2 and #5 at 5 d. **c**, Relative leaf water content of WT, *GC1∷GAD2Δ* #2 and #5 plants following drought treatment for 0 d, 5 d, 7 d and 10 d; n = 6 for 0 d, 5 d and 5 d samples, except n = 18 for WT at 10 d, n = 12 for *GC1∷GAD2Δ* #2 at 10 d and n = 13 for *GC1∷GAD2Δ* #5 at 10 d. **d**, Representative images of WT, *GC1∷GAD2Δ* #2 and #5 plants before (0 day) and after (10 days) drought treatment as indicated. Pot size 2.5 inch diameter x 2.25 inch height (LiCor). All data are plotted (**b**) or represented as mean ± s.e.m (**c**); statistical difference was determined using One-way ANOVA (**b**) or Two-way ANOVA (**c**); \**P*< 0.05 and \*\*\**P*< 0.001.

### GABA negatively regulates activity of ALMTs in guard cells

ALMTs are plant-specific anion channels that share no homology to Cys-loop receptors except a region of 12 amino acid residues predicted to bind GABA in GABA_A_ receptors^13, 22^. In animals, ionotropic GABA receptors are stimulated by GABA; in contrast, anion currents through ALMTs are inhibited by GABA^10, 11^. To investigate whether ALMTs constitute the mechanism by which GABA signals are transduced in plants, we examined the GABA sensitivity of T-DNA insertional mutants that lack expression of two guard cell localised Arabidopsis ALMT proteins.

ALMT9 is the dominant tonoplast-localised channel involved in anion uptake into guard cell vacuoles during stomatal opening, but has no documented role in closure^15^. We hypothesised that GABA might target and inhibit ALMT9 activity to reduce the rate or extent of stomatal opening. To test this we crossed two *almt9* alleles (*almt9-1* and *almt9-2*) with *gad2-1*. We found that, similar to *gad2*, both double mutants (*almt9-1/gad2-1* and *almt9-2/gad2-1*) maintained low GABA accumulation in their leaves (Fig. 6; Supplementary Fig. 9a-d). However, unlike *gad2*, both *almt9-1/gad2-1* and *almt9-2/gad2-1* had wildtype like stomatal conductance (Fig. 6; Supplementary Fig. 9a-d). We also tested the GABA sensitivity of single *almt9* mutants. We found, in contrast to wildtype, that light-induced stomatal opening in *almt9* lines was not sensitive to exogenous GABA or muscimol (Fig. 7a, b; Supplementary Fig. 9e, f), whereas dark-induced stomatal closure in *amlt9* retained in its GABA sensitivity (Fig. 7c, d ; Supplementary Fig. 9g, h). These results are consistent with GABA reducing stomatal opening via negative regulation AMLT9-mediated Cl^−^ uptake into guard-cell vacuoles. Furthermore, it strongly indicates the corollary of this finding, that the higher stomatal conductance phenotype of *gad2* is the result of greater ALMT9 activity due to its lack of inhibition by GABA.

**Figure 6.**
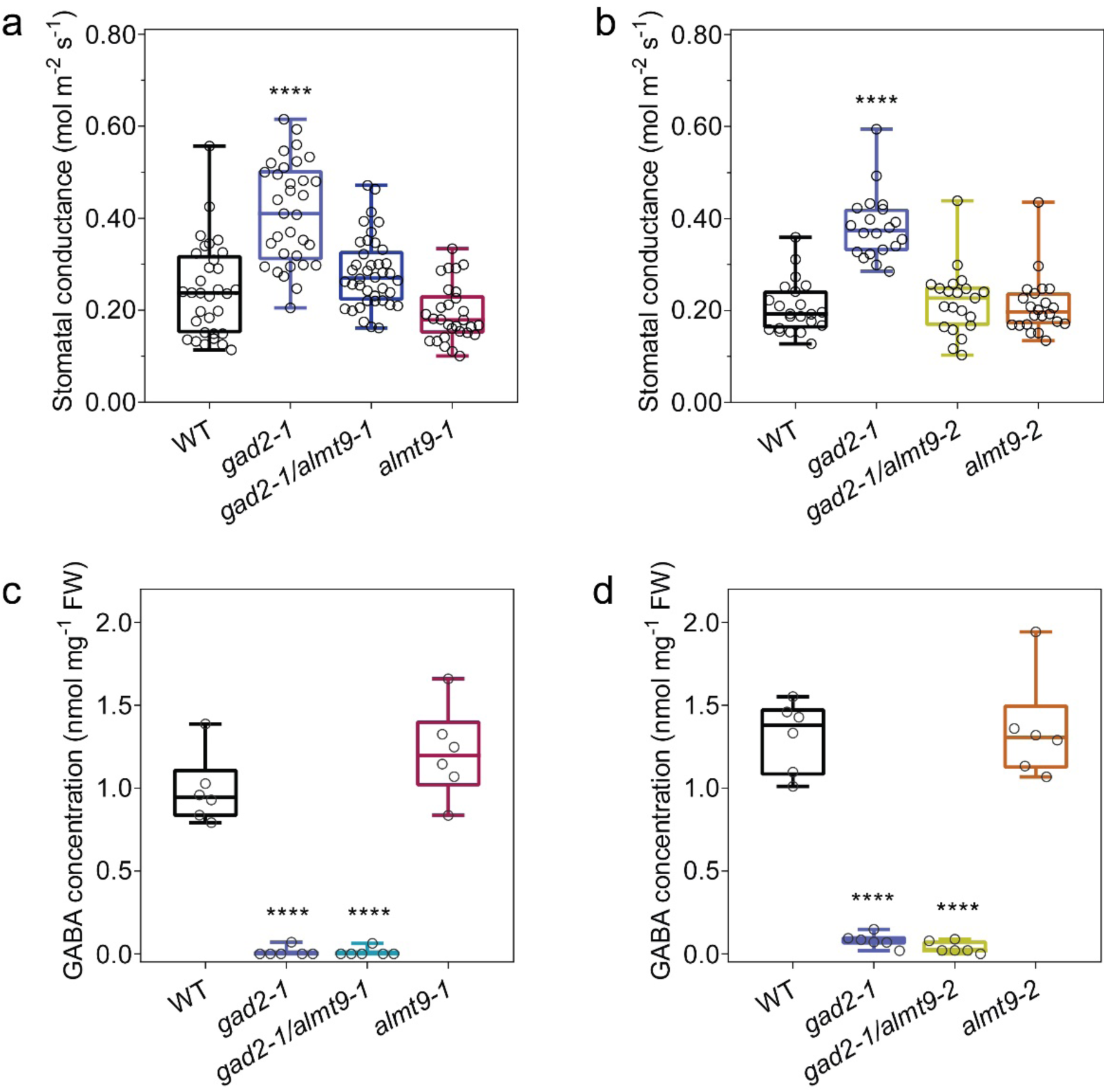
The loss of ALMT9 attenuates the *gad2* mutant phenotype and abolishes GABA suppression of stomatal opening. **a-d**, Leaf stomatal conductance (**a, b**) and GABA concentration (**c, d)** of WT, *gad2-1, gad2-1/almt9-1, almt9-1, gad2-1/almt9-2* and *almt9-2* plants; n = 32 for WT, n = 33 for *gad2-1*, n = 40 for *gad2-1/almt9-1* and n = 29 for *almt9-1*, data collected from three independent batches of plants (**a**); n = 22 for WT, n = 20 for *gad2-1*, n = 21 for *gad2-1/almt9-2* and n = 22 for *almt9-2*, data collected from two independent batches of plants (**b**); n = 6 plants (**c, d**). All data are plotted, statistical difference was determined by One-way ANOVA (**a-d**), \**P*<0.05, \*\*\**P*<0.001 and \*\*\*\**P*<0.0001.

**Figure 7.**
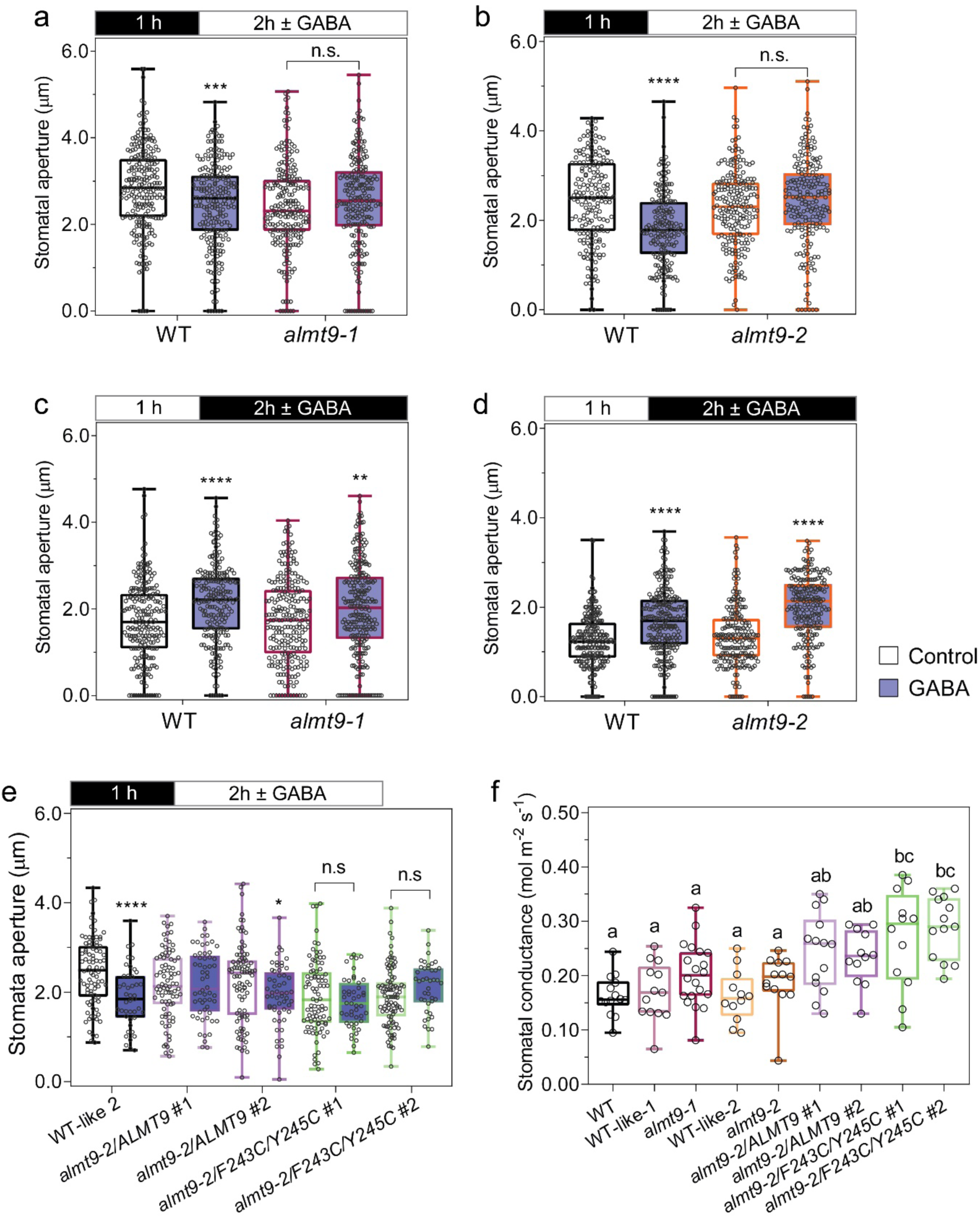
GABA sensitivity of stomatal opening is lost in *almt9* and restored by *ALMT9* complementation but not *ALMT9*^F243C/Y245C^. **a-e**, Wildtype, a*lmt9* knockout and complementation plant stomatal aperture in response to dark or light. Epidermal strips were pre-incubated in stomatal measurement buffer for 1 h under dark (**a, b, e**) or light (**c, d**), followed by 2 h in light (**a, b, e**) or dark (**c, d**) as indicated by black (dark) or white (light) bars above graphs, ± 2 mM GABA; n = 236 for WT and n = 221 for *almt9-1* with control treatment, n = 229 for WT and n = 215 for *almt9-1* with GABA treatment (**a**); n = 223 for WT and n = 242 for *almt9-1* with control treatment, n = 215 for WT and n = 256 for *almt9-1* with GABA treatment (**b**); n = 183 for WT and n = 189 for *almt9-2* with control treatment, n = 210 for WT and n = 197 for *almt9-2* with GABA treatment (**c**); n = 236 for WT and n = 243 for *almt9-2* with control treatment, n = 202 for WT and n = 220 for *almt9-2* with GABA treatment (**d**); n = 88 for WT-like-2 (segregated from *almt9-2*)^15^, n = 85 for *almt9-2* complement with *35S∷ALMT9* #1 (*almt9-2/ALMT9* #1), n = 93 for *almt9-2/ALMT9* #2, n = 88 for *almt9-2* complement with *35S∷ALMT9* with double mutation F243C/Y245C as putative GABA interaction residues^13, 22^ (*almt9-2/*F243C/Y245C #1), and n = 98 for *almt9-2/*F243C/Y245C #2 with control treatment; n = 43 for WT-like-2, n = 56 for *almt9-2/ALMT9* #1 and #2, n = 47 for *almt9-2/*F243C/Y245C #1, and n = 41 for *almt9-2/*F243C/Y245C #2 with GABA treatment (**e**). **f**, Stomatal conductance of wildtype, a*lmt9* knockout and complementation plants; n = 14 for WT and *almt9-2/ALMT9* #1, n = 13 for WT-like-1 (segregated from *almt9-1*)^15^, *almt9-2* and *almt9-2/*F243C/Y245C #2, n = 20 for *almt9-1*, n = 12 for WT-like-2, *almt9-2/ALMT9* #2 and *almt9-2/*F243C/Y245C #1. All data are plotted, statistical difference was determined by Two-way ANOVA (**a-d**), Student’s *t-*test (**e**) or One-way ANOVA (**f**), \**P*<0.05, \*\**P* <0.01, \*\*\**P* <0.001 and \*\*\*\**P* <0.0001 (**a-e**); a, b, c represent groups with no significant difference, *P*<*0*.*01* (**f**).

We further tested this hypothesis by attempting to complement *almt9* plants with either the native channel or a site-directed ALMT9 mutant (ALMT9^F243C/Y245C^). The mutations within ALMT9^F243C/Y245C^ are in the 12 amino acid residue motif that shares homology with a GABA binding region in mammalian GABA_A_ receptors^13,22^. Mutations in the aromatic amino acid residues in this motif have been shown for other ALMTs to result in active channels that are not inhibited by GABA when tested in heterologous systems^30^. However, no *in planta* tests of whether these mutations in the GABA binding site of ALMT result in a transport competent protein that lacks GABA sensitivity have been performed to date. Here, we found that - similar to *almt9* lines - stomatal opening of *almt9-2* expressing *ALMT9*^*F243C/Y245C*^ were insensitive to GABA, unlike wildtype plants and native ALMT9 expression line 2 (Fig. 7e). Furthermore, the steady state stomatal conductance of both *ALMT9*^*F243C/Y245C*^ lines were significantly greater than that of wildtype and *almt9* lines (Fig. 7f). This result indicates that we successfully complemented *almt9* with an active but GABA-insensitive form of ALMT9, and that this increased transpirational water loss over wildtype levels. These data are completely consistent with ALMT9 being a GABA target, and through this process GABA regulating plant water loss, even under non-stressed conditions. The effect of GABA is then amplified under a water deficit when its concentration increases. We propose that GABA accumulation has a role in promoting drought tolerance by reducing the amplitude of stomatal re-opening each morning, which minimises whole plant water loss. As such, the GABA-ALMT interaction is a strong candidate for constituting the ABA-independent stress memory of a decreased soil water status that has been previously proposed without mechanistic attribution^31, 32^.

Next we examined plasma membrane-localised ALMT12, which is involved in stomatal closure but not opening^14^ (Fig. 1). We observed that, unlike wildtype plants, closure of *almt12* knockouts were insensitive to GABA or muscimol when transitioning from light-to-dark (Fig. 8a; Supplementary Fig. 10a). In contrast, stomatal opening of *almt12* lines showed wildtype-like sensitivity to GABA or muscimol when transitioning from dark-to-light (Fig. 8b; Supplementary Fig. 10b). These data indicate that ALMT12 is a plasma membrane GABA target that affects stomatal closure in response to dark. Guard cells express additional ALMTs^16, 33^; however, the stomatal pore apertures of *almt9 x almt12* double knockouts exhibited no sensitivity to GABA or muscimol following light and dark transitions (Fig. 8c, d; Supplementary Fig. 10c, d). This indicates that ALMT12 and ALMT9 are the major GABA targets modulating stomatal movement, and that additional experiments are required to resolve whether other ALMTs further contribute to GABA-regulation of stomata (Fig. 9).

**Figure 8.**
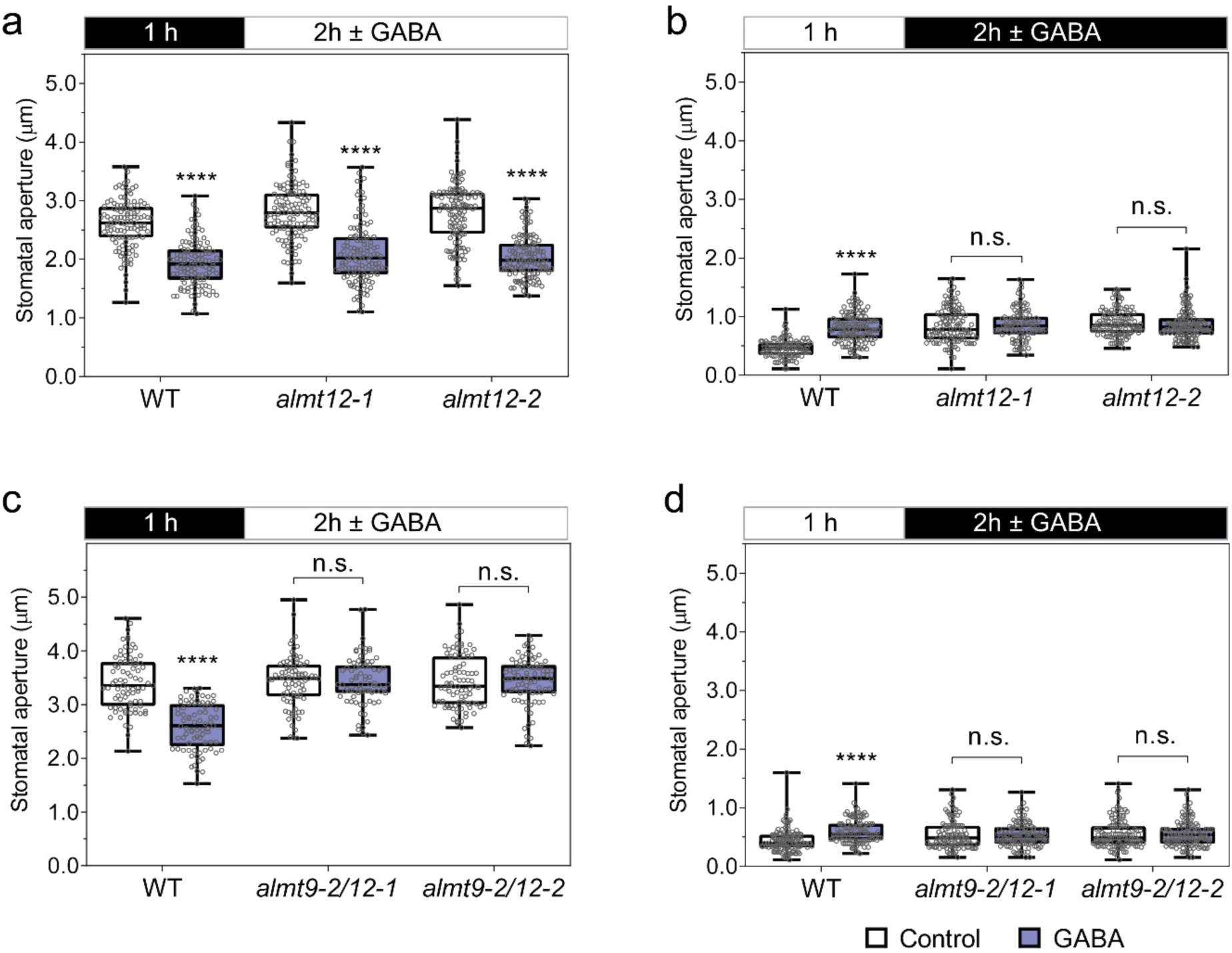
The loss of ALMT12 abolishes GABA suppression of stomatal closure. **a-b**, Stomatal aperture measurement of wildtype (WT), *almt12* and *almt9-2/almt12* knockout plants in response to dark or light. Epidermal strips were pre-incubated in stomatal measurement buffer for 1 h under dark (**a**) or light (**b**), followed by 2 h incubation in light (**a, c**) or dark (**b, d**) as indicated by black (dark) or white (light) bars above the plots in absence or presence of 2 mM GABA; n = 116 for WT (control), n = 119 for *almt12-1* (control), n = 120 for *almt12-2* (control), n = 113 for WT (GABA), n = 123 for *almt12-1* (GABA), n = 124 for *almt12-2* (GABA) (**a**); n = 105 for WT (control), n = 115 for *almt12-1* (control), n = 122 for *almt12-2* (control), n = 122 for WT (GABA), n = 100 for *almt12-1* (GABA), n = 131 for *almt12-2* (GABA) (**b**); n = 78 for WT (control), n = 82 for *almt9-2/12-1* (control), n = 83 for *almt9-2/12-2* (control), n = 77 for WT (GABA), n = 77 for *almt9-2/12-1* (GABA) and n = 81 for *almt9-2/12-2* (GABA) (**c**); n = 114 for WT (control), n = 104 for *almt9-2/12-1* (control), n = 120 for *almt9-2/12-2* (control), n = 113 for WT (GABA), n = 114 for *almt9-2/12-1* (GABA), n = 123 for *almt9-2/12-2* (GABA) (**d**). All data are plotted, statistical difference was determined using Two-way ANOVA, \*\*\*\**P* < 0.0001; all experiments were repeated at least twice from different batches of plants with blind treatments (**a-d**).

**Figure 9.**
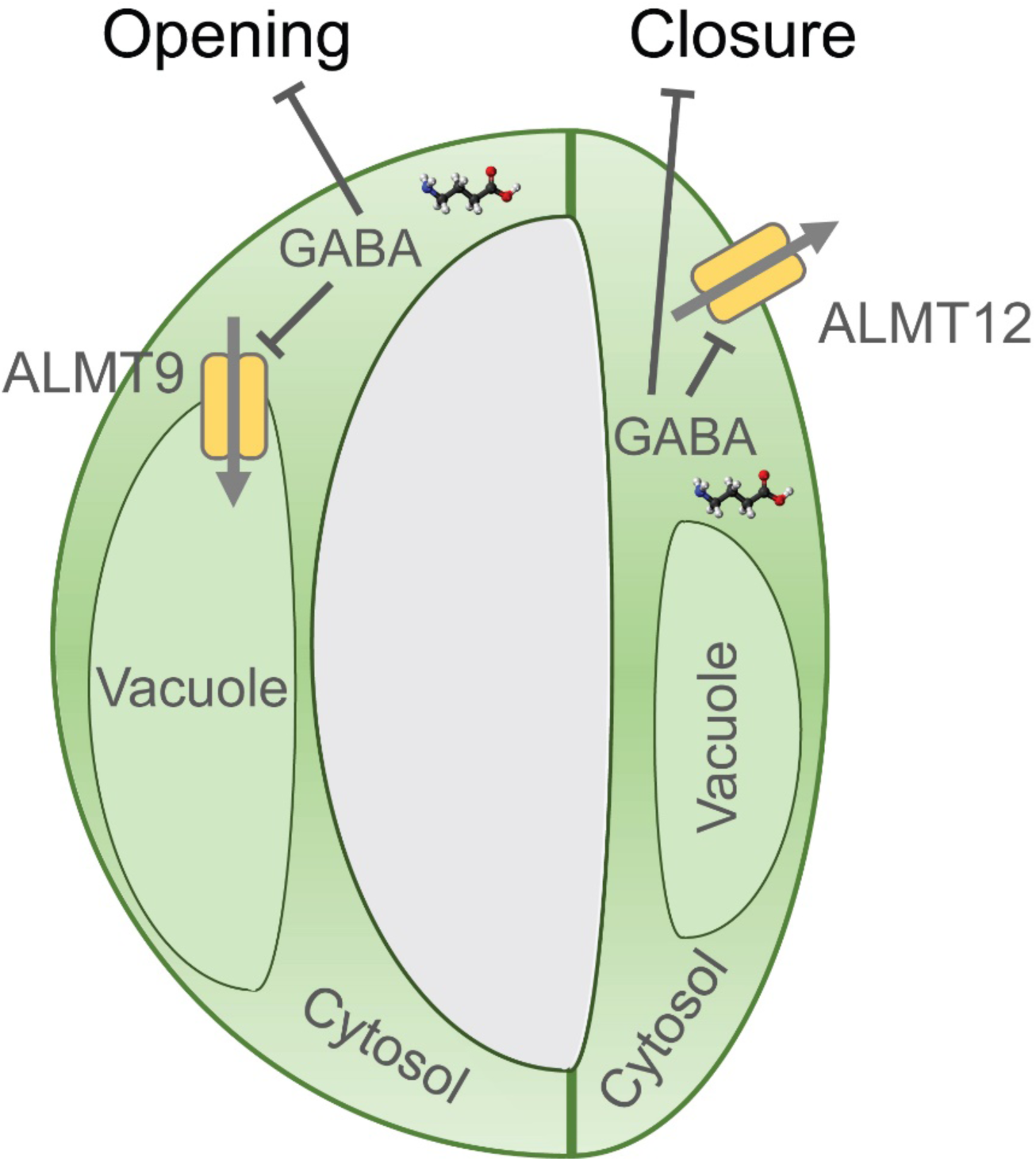
Proposed model of how GABA-mediated signaling regulates stomatal movement. Cytosolic guard cell GABA antagonizes stomatal movement via negative regulation of ALMT9 and ALMT12, which respectively control stomatal opening and closure. The site of primary action of endogenous cytosolic GABA is ALMT9, reducing the extent of opening. When GABA increases under stress, GABA, rather than stimulating movement reduces opening the subsequent day. This constitutes a ‘memory’ of stress whilst GABA levels are maintained above baseline. The physiological role of GABA block of ALMT12 activity is still to be ascertained.

## Discussion

GABA concentration oscillates over diel cycles and increases in response to multiple abiotic and biotic stresses including drought, heat, cold, anoxia, wounding pathogen infection and salinity^23^. ALMT have been implicated in modulating multiple developmental and physiological processes in plants^14, 15, 16, 34, 35, 36^ including those underpinning nutrient uptake and fertilization that are affected by GABA^13, 37^. Therefore, our discovery that GABA regulates ALMT to form a novel and physiologically relevant signalling mechanism in guard cells is likely to have broad significance beyond stomata, particularly during plant responses to environmental transitions and stress. GABA’s effect on stomata appears to be conserved across a large range of crops from diverse clades including important monocot and dicot crops, indicating that GABA is stomatal signal of economic significance. As we find that the genetic manipulation of cell-type specific GABA metabolism can reduce water loss and improve drought performance our work opens up a new frontier of research for manipulating crop stress resilience. Furthermore, as we provide the first conclusive proof that GABA is a plant signalling molecule and not just a plant metabolite^12, 18^, we definitively demonstrate that GABA is an endogenous signalling molecule beyond the animal and bacterial kingdoms, enacted through distinct and organism specific mechanisms.

## METHODS

### Plant materials and growth conditions

All experiments performed on *Arabidopsis thaliana* were in the Columbia-0 (Col-0) ecotype background, unless stated. Arabidopsis wildtype, T-DNA insertion mutant and other transgenic plants were germinated and grown on ½ Murashige and Skoog (MS) medium with 0.8% phytagel for 10 days before being transferred to soil for growth in short-day conditions (100-120 µmol m^-2^ s^-1^, 10 h light/14 h dark) at 22 °C. The T-DNA insertion mutant *gad2-1* (GABI_474_E05) and *gad2-2* (SALK_028819) were obtained from the Arabidopsis Biological Resource Centre (ABRC). *gad2-1* was selected using primer sets: gad2_LP1 (5’-TATCACGCTAACACCTAACGC-3’), gad2_RP1 (5’-TTCAAGGTTTGTCGGTATTGG-3’), and, GABI_LB (5’-GGGCTACACTGAATTGGTAGCTC-3’) for removing the second T-DNA insert; gad2_LP2 (5’-ACGTGATGGATCCAGACAAAG-3’), gad2_RP2 (5’-TCTTCATTTCCACACAAAGGC-3’), and, GABI_LB for isolation of the *GAD2* (At1g65960) T-DNA insertion. *gad2-2* was selected using primer sets: gad2-2_LP (5’-AGTTGTATGAAAGTTCATGTGGC-3’), gad2-2_RP (5’-TCGACCACGAGATTTTAATGG -3’), and, SALK_LB (5’-ATTTTGCCGATTTCGGAAC-3’). *almt9-1* (SALK_055490), *almt9-2* (WiscDsLox499H09), *almt12-1* (SM_3_38592) and *almt12-2* (SM_3_1713) were selected as described previously^14,15^. The double mutant lines *gad2-1/almt9-1, gad2-1/almt9-2, almt9-2/12-1* and *almt9-2/12-2* were obtained respectively from crossing the respective mutants. The mesophyll enhancer-trap line JR11-2 in the Columbia-0 background was kindly provided by K. Baerenfaller (ETH Zurich)^38^. JR11-2(Col-0) and *gad2-1*/JR11-2 were segregated from crossing *gad2-1* with JR11-2. JR11-2 was selected using primer sets: JR11-2_LP (5’-TTATTTAGGGAAATTACAAGTTGC-3’), JR11-2_RP (5’-AGACACATTTAATAACATTACAACAAA-3’) and JR11-2_LB (5’-GTTGTCTAAGCGTCAATTTGTTT-3’)^39^. All experiments were performed on stable T_3_ transgenic plants or confirmed homozygous mutant lines. The other plants –*Vicia faba, Nicotiana benthamiana* and *Glycine max* were grown in soil in long-day conditions (400 µmol m^-2^ s^-1^, 16 h light/8 h dark, 28 °C/25 °C). *Hordeum vulgare* (barley) cv. Barke was grown in a hydroponic system with half-strength Hoagland’s solution in long-day conditions (150 µmol m^-2^ s^-1^, 16 h light/8 h dark, 23 °C).

### Gene cloning and plasmid construction

For guard-cell specific complementation, the constitutively active form of *GAD2* with a truncation of calmodulin binding domain (*GAD2Δ*)^26, 28^ and the full-length *GAD2* coding sequence (*GAD2*) was driven by a guard-cell specific promoter *GC1* (−1140/+23)^27^, as designated *GC1∷GAD2Δ* and *GC1∷GAD2* respectively. PCR reactions first amplified the truncated *GAD2Δ* with a stop codon and *GC1* promoter (*GC1*) separately using Phusion^®^ High-Fidelity DNA Polymerase (New England Biolabs) with the primer sets: GAD2_forward (5’-CACTACTCAAGAAATATGGTTTTGACAAAAACCGC-3’) and GAD2_truncated_reverse (5’-TTATACATTTTCCGCGATCCC-3’); GC1_forward (5’-CACCATGGTTGCAACAGAGAGGATG-3’) and GC1_reverse (5’-ATTTCTTGAGTAGTGATTTTGAAG-3”). This was followed by an overlap PCR to fuse the *GC1* promoter to *GAD2Δ* (*GC1∷GAD2Δ*) with the GC1_forward and GAD2_truncated_reverse primer set. The same strategy was used to amplify *GC1∷GAD2Δ* without a stop codon (*GC1∷GAD2Δ-stop*), *GC1∷GAD2* and *GC1∷GAD2* without a stop codon (*GC1∷GAD2-stop*) with different primer sets: 1) *GC1∷GAD2Δ-stop* amplified with GAD2_forward and GAD2_truncated-stop_reverse (5’-TACATTTTCCGCGATCCCT-3’); 2) *GC1∷GAD2* amplified with GAD2_forward and GAD2_reverse (5’-TTAGCACACACCATTCATCTTCTT-3’); 3) *GC1∷GAD2-stop* amplified with GAD2_forward and GAD2-stop_reverse (5’-CACACCATTCATCTTCTTCC-3’). The fused PCR products were cloned into the pENTR/D-TOPO vector via directional cloning (Invitrogen). pENTR/D-TOPO vectors containing *GC1∷GAD2Δ* or *GC1∷GAD2* were recombined into a binary vector pMDC99^40^ by a LR reaction using LR Clonase II Enzyme mix (Invitrogen) for guard-cell specific complementation, after an insertion of a NOS Terminator into this vector. A pMDC99 vector was cut by *Pac*I (New England Biolabs) and ligated with NOS terminator flanked with *Pac*I site using T4 DNA ligase (New England Biolabs). This NOS terminator flanked with *Pac*I site was amplified with primer set: nos_PacI_forward (5’-TACGTTAATTAAGAATTTCCCCGAT-3’) and nos_PacI_reverse (5’-GCATTTAATTAAAGTAACATAGATGACACC-3’) and cut by restriction enzyme *Pac*I before T4 DNA ligation. *GC1∷GAD2Δ-stop* and *GC1∷GAD2-stop* were recombined from the pENTR/D-TOPO vector into a pMDC107 vector that contained a GFP tag on the C-terminus (*GC1∷GAD2Δ-GFP* and *GC1*∷ *GAD2-GFP*)^40^.

To create *GAD2* complementation driven by a constitutive 35S promoter, the full-length *GAD2* was also amplified using primer set GAD2_forward2 (5’-CACCATGGTTTTGACAAAAACCGC-3’) and GAD2_reverse and cloned into pENTR/D-TOPO vector via directional cloning (Invitrogen), followed by a LR reaction recombinant into pMDC32^40^. For mesophyll specific complementation, *GAD2Δ* with a stop codon was amplified with the GAD2_forward2 and GAD2_truncated_reverse primer set, and cloned into the pENTR/D-TOPO vector, followed by a LR reaction recombined into the pTOOL5 vector (*UAS∷GAD2Δ*)^41^.

For *GAD2* expression analysis, a 1 kb sequence upstream of the *GAD2* start codon was designated as the *GAD2* promoter (*pGAD2*) and amplified using primer set proGAD2_F (5’-ATTTTGAATTTGCGGAGAATCT-3’) and proGAD2_R (5’-CTTTGTTTCTGTTTAGTGAAAGAGAA-3’). The *pGAD2* PCR product was cloned into pCR8/GW/TOPO via TA cloning and recombined via a LR reaction into the pMDC162 vector containing the *GUS* reporter gene for histochemical assays^40^. The binary vectors, pMDC32, pMDC99, pMDC107, pMDC162 and pTOOL5 carrying sequence-verified constructs were transformed into *Agrobacterium* strain *AGL1* for stable transformation in Arabidopsis plants.

### Stomatal aperture and density measurement

Soil-grown Arabidopsis (5-6 week-old) were used for stomatal aperture and density measurements. Two to three week-old soybean, broad beans and barley, and 5-6 week-old tobacco were used for stomatal aperture assays. Epidermal strips from Arabidopsis, soybean, faba bean and tobacco were peeled from abaxial sides of leaves, pre-incubated in stomatal measurement buffer containing 10 mM KCl, 5 mM L-malic acid, 10 mM 2-ethanesulfonic acid (MES) with pH 6.0 by 2-Amino-2-(hydroxymethyl)-1,3-propanediol (Tris) under light (200 µmol m^-2^ s^-1^) or darkness, and transferred into stomatal measurement buffer with blind treatments as stated in the figure legend. For barley epidermal stomatal assays, a modified method was used^42^: the second fully expanded leaf from 2 week-old seedlings was used as experimental material, leaf samples were first detached and bathed in a modified measurement buffer (50 mM KCl, 10 mM MES with pH 6.1 by KOH) under light (150 µmol m^-2^ s^-1^) or darkness for 2 h, then pre-treated in the same buffer with or without 1 mM GABA for 0.5 h; after this pre-treatment, samples were incubated in continuous dark, light, light-to-dark or dark-to-light transition for an additional 1 h as indicated in the figure legend before leaf epidermal strips were peeled for imaging. For Arabidopsis stomatal density measurement, epidermal strips were peeled from abaxial sides of young and mature leaves, three leaves per plants, three plants per genotype. Epidermal strips for both aperture and density measurement were imaged using an Axiophot Pol Photomicroscope (Carl Zeiss) apart from the barley epidermal strips imaged using an Nikon Diaphot 200 Inverted Phase Contrast Microscope (Nikon). Stomatal aperture and density were analyzed using particle analysis (http://rsbweb.nih.gov/ij/).

### Stomatal conductance measurement

All stomatal conductance measurements were performed on 5-6 week-old Arabidopsis plants. The stomatal conductance determined by the AP4 Porometer (Delta-T Devices) was calculated based on the mean value from two to three leaf recordings per plant (Fig. 3b, 4c, 5b, 6a, b, 7f; Supplementary Fig. 7d, h, j). The stomatal conductance was also recorded using LI-6400XT Portable Photosynthesis System (LI-COR Biosciences) equipped with an Arabidopsis leaf chamber fluorometer (under 150 µmol m^-2^ s^-1^ light with 10% blue light, 150 mmol s^-1^ flow rate, 400 ppm CO_2_ mixer, ∼50 % relative humidity at 22 °C) as indicated (Fig. 4d, e). The stomatal conductance of the detached leaf assay was measured using LCpro-SD Portable Photosynthesis System (ADC Bioscientific) with 350 µmol m^-2^ s^-1^ light, 200 µmols s^-1^ flow rate and 400 ppm CO_2_ at 22 °C (Supplementary Fig. 1c-e). The detached leaf was fed with artificial xylem sap solution modified as described^43^, containing 1 mM KH_2_PO_4_, 1 mM K_2_HPO_4_, 1 mM CaCl_2_, 0.1 mM MgSO_4_, 1 mM KNO_3_, 0.1 mM MnSO_4_, 1 mM K-H-malate, pH 6.0 (KOH).

### GABA measurement

GABA concentration was determined using Ultra Performance Liquid Chromatography (UPLC) as described previously^30^. Briefly, GABA was extracted from samples using 10 mM sodium acetate and derivatized with the AccQ Tag Ultra Derivatization Kit (Waters). Chromatographic analysis of GABA was performed on an Acquity UPLC System (Waters) with a Cortecs or Phenomenex UPLC C18 column (1.6 μm, 2.1×100 mm). The gradient protocol for amino acids analysis was used to measure GABA with mobile solvents AccQ Tag Ultra Eluents A and B (Waters). Standard GABA solution was used for calibration ranged from 0 to 150 µM. The results were analyzed by Empower chromatography software (Waters).

### GUS histochemical staining assays

A GUS histochemical assay was performed using the methods described previously^44^. 3-4 week-old transgenic *pGAD2∷GUS* plants were stained in buffer containing 50 mM Na phosphate pH = 7.0, 10 mM EDTA, 2 mM potassium ferrocyanide, 2 mM potassium ferricyanide, 0.1% (v/v) Triton X-100, 0.1% (w/v) X-Gluc (5-bromo-4-chloro-3-indolyl β-d-glucuronide) during a 1.5 h incubation at 37 °C in the dark. The stained plants were destained in 70% ethanol. GUS-stained plants were imaged using an Axiophot Pol Photomicroscope (Carl Zeiss).

### Leaf relative water content and soil water measurement

Fresh weight of leaves was determined immediately after abscission on an Ohaus Explorer E02140 Balance, then sampled leaves were rehydrated to full turgid weight in ultrapure water overnight and measured after surface water was dried with paper towel. Dry weight was determined at 65 °C for 1 d. Leaf relative water content (RWC) was calculated by an equation:

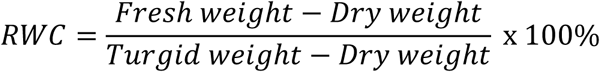

Fresh soil weight of the whole pot (Mwet) and dry soil weight after drying the soil (Mdry) at 105 °C for 3 d was measured using an Ohaus ARA520 Adventurer Balance. Gravimetric soil water content (θg) of the whole soil in the pots was calculated by an equation:

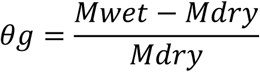

### Fluorescence microscopy

The fluorescence of fluorescent proteins in transgenic *gad2-1/GC1∷GAD2Δ-GFP* and *gad2-1/GC1∷GAD2-GFP* plants was imaged by confocal laser scanning microscopy using a Zeiss Axioskop 2 mot plus LSM5 PASCAL and argon laser (Carl Zeiss). Sequential scanning and laser excitation was used to capture fluorescence via the LSM5 PASCAL from GFP (excitation = 488 nm, emission Band-Pass = 505-530 nm), chlorophyll autofluorescence (excitation = 543 nm, emission Long-Pass = 560 nm).

### Reverse transcriptional PCR

Reverse transcriptional PCR was determined by PCR amplification on cDNA synthesized from RNA extracted from plants as indicated. PCR amplified *GAD2, Actin2, GFP, GAD2 mRNA, UAS∷GAD2Δ* and *ALMT9* using Phire Hot Start II DNA Polymerase (Invitrogen) with primer sets:

GAD2_rt_F (5’-ACGTGATGGATCCAGACAAAG-3’) and

GAD2_rt-R (5’-TACATTTTCCGCGATCCCT-3’);

Actin2_rt_F (5’-CAAAGGCCAACAGAGAGAAGA-3’) and

Actin2_rt_R (5’-CTGTACTTCCTTTCAGGTGGTG-3’);

GFP_rt_F (5’-GGAGTTGTCCCAATTCTTGTT-3’) and

GFP_rt_R (5’-CGCCAATTGGAGTATTTTGT-3’);

GAD2mRNA_rt_F (5’-ACGTGATGGATCCAGACAAAG-3’) and

GAD2mRNA_rt_R (5’-TCTTCATTTCCACACAAAGGC-3’);

UAS_GAD2_rt_F (5’-TCACTCTCAATTTCTCCAAGG-3’) and

UAS_GAD2_rt_R (5’-CGGCAACAGGATTCAATCTTAAG-3’);

ALMT9_rt_F (5’-AATACTCGAGAAACGGGAGAG-3’) and

ALMT9_rt_R (5’-CATCCCAAAACACCTACGAAT-3’).

### Quantitative real-time PCR analysis

Quantitative reverse transcription PCR was performed using KAPA SYBR FAST ABI PRISM kit (Kapa Biosystems) using a QuantStudio™ 12K Flex Real-Time PCR System (Thermo Fisher Scientific) to determine the expression levels of *GAD1, GAD2, GAD3, GAD4, GAD5, GABA-T, ALMT9, ALMT12, RD29A* and *RD22* genes with primer sets:

GAD1_qF (5’-TCTCAAAGGACGAGGGAGTG-3’) and

GAD1_qR (5’-AACCACACGAAGAACAGTGATG-3’);

GAD2_qF (5’-GTCTCAAAGGACCAAGGAGTG-3’) and

GAD2_qR (5’-CATCGGCAGGCATAGTGTAA-3’);

GAD3_qF (5’-CCGTTAGTGGCGTTTTCTCT-3’) and

GAD3_qR (5’-TCTCTTTGCGTCTCCTCTGG-3’);

GAD4_qF (5’-GTGTTCCGTTAGTGGCGTT-3’) and

GAD4_qR (5’GTCTCCTCTGGCGTCTTCTT-3’);

GAD5_qF (5’-TCAACCCACTTTCACTCTCA-3’) and

GAD5_qR (5’-TTCCTTCTCTTAGCCTCCTT-3’);

GABA-T_qF (5’-AGGCAGCACCTGAGAAGAAA-3’) and

GABA-T_qR (5’-GGAGTGATAAAACGGCAAGG-3’);

ALMT9_qF (5’-CAGAGAGTGGGCGTAGAAGG-3’) and

ALMT9_qR (5’-GGATTTGAAGGCGTAGATTGG-3’);

ALMT12_qF (5’-TTGACGGAACTCGCAGATAG-3’) and

ALMT12_qR (5’-CGATGGAGGTTAGAGCCAAG-3’);

RD29A_qF (5’-AAACGACGACAAAGGAAGTG-3’) and

RD29A_qR (5’-ACCAAACCAGCCAGATGATT-3’);

RD22_qF (5’-AGGGCTGTTTCCACTGAGG-3’) and

RD22_qR (5’-CACCACAGATTTATCGTCAGACA-3’).

Expression levels of each gene was normalised to three control genes – *Actin2, EF1α* and *GAPDH-A* that were amplified with primer sets:

Actin2_qF (5’-TGAGCAAAGAAATCACAGCACT-3’) and

Actin2_qR (5’-CCTGGACCTGCCTCATCATAC-3’);

EF1α_qF (5’-GACAGGCGTTCTGGTAAGGAG-3’) and

EF1α_qR (5’-GCGGAAAGAGTTTTGATGTTCA-3’);

GAPDH-A_qF (5’-TGGTTGATCTCGTTGTGCAGGTCTC-3’) and

GAPDH-A_qR (5’-GTCAGCCAAGTCAACAACTCTCTG-3’).

## Data availability

Source data for Figs. 1-9 and Supplementary Figs. 1-10 are provided with the paper. Sequence data used in this paper can be found in The Arabidopsis Information Resource (TAIR) database (https://www.arabidopsis.org/) under the following accessions: *GAD1* (At5g17330), *GAD2* (At1g65960), *GAD3* (At2g02000), *GAD4* (At2g02010), *GAD5* (At3g17760), *GABA-T* (At3g22200), *ALMT9* (At3g18440), *ALMT12* (At4g17970), *RD29A* (At5g52310) and *RD22* (At5g25610). Other data that support the findings of this study are available from the corresponding author upon request.

## Acknowledgements

We would like to thank Dr. Katja Bärenfaller from University of Zurich for providing JR11-2 (Col-0) seeds; A/Prof. Zhonghua Chen from Western Sydney University for assisting with barley epidermal assays; Dr. Cornelia Eisenach from University of Zurich for providing *ALMT9* constructs, *almt9* and *almt12* seeds; Prof. Stephen D Tyerman, Prof. Enrico Martinoia and Dr Alexis De Angeli for valuable discussions.

## Author contributions

BX constructed all materials and performed all experiments except the following: YL generated and performed experiments on *almt12* and *almt9xalmt12* materials. XZ performed all non-Arabidopsis aperture measurements, except for barley performed by NS. LC performed GABA measurements supervised by MO. JH acquired images used in Fig 1. MG, BX and RH conceived the research. BX drafted all figures. MG and BX drafted the manuscript, all authors provided edits. This work was funded by Waite Research Institute Funding to MG and BX, ARC Discovery grant DP170104384 to MG and RH, ARC Centre of Excellence (CE1400008) and Grains Research and Development Corporation funding (UWA00173) to MG. XZ was supported by a Chinese Scholarship Council Travelling Fellowship.

## Additional Information

Extended Data is available for this paper.

