## Supplemental Data for "GABA signalling in guard cells acts as a ‘stress memory’ to optimise plant water loss"

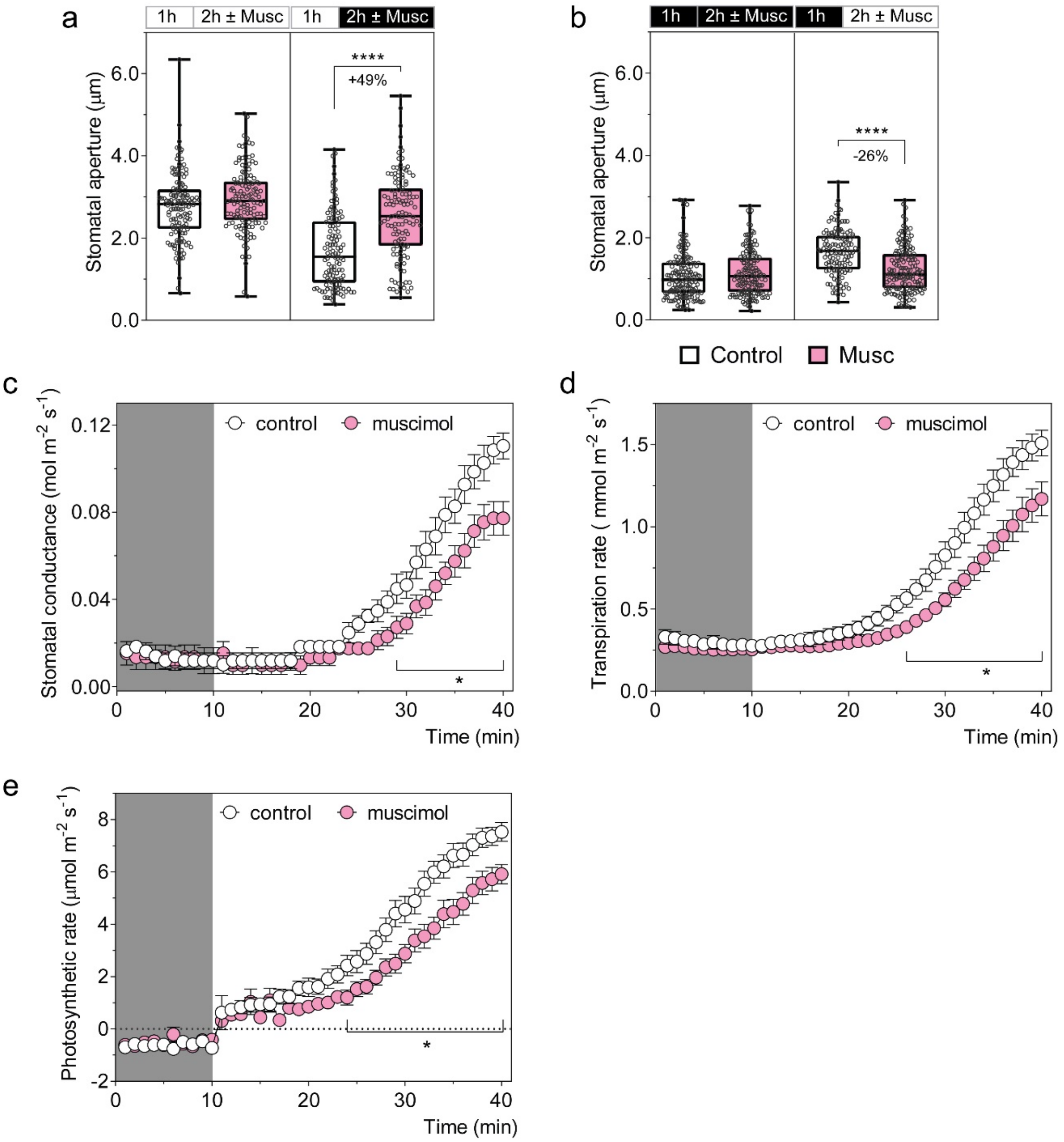


**Supplementary Figure 1. Muscimol antagonises stomatal movement initiated by light and dark treatments. a-b,** Exogenous muscimol application reduces stomatal movement in response to light or dark. Epidermal strips were pre-incubated in stomatal measurement buffer for 1 h under light (**a**) or dark (**b**), followed by 2 h incubation under constant light (**a**), dark (**b**), light-to-dark transition (**a**) or dark-to-light transition (**b**) as indicated above graphs by black (dark) or white (light) bars, together with the application of 10 µM muscimol (Musc); n = 134 for control (constant light), n = 132 for muscimol (constant light), n = 118 for control (light-to-dark transition) and n = 120 for muscimol (light-to-dark transition) (**a**); n = 156 for control (constant dark), n = 151 for muscimol (constant dark), n = 127 for control (dark-to-light transition) and n = 151 for muscimol (dark to light transition) (**b**). All data are plotted (a-b); statistical difference was determined using Two-way ANOVA, *****P*< 0.0001. **c**, Stomatal conductance of detached leaves of wildtype Arabidopsis plants determined by LCpro-SD Portable Photosynthesis System. A detached leaf was first fed artificial xylem sap solution with or without 10 μM muscimol for 30 min under light to allow uptake of treatment, followed by 30 min in the dark to close stomata before recording; data represent mean ± s.e.m, n = 7 for control and n = 8 for muscimol; statistical difference was determined by Student's *t-test*, **P< 0.05*. **d-e**, Transpiration rate (**d**) and photosynthesis rate (**e**) of detached leaves fed artificial xylem sap solution recorded by a LCpro-SD Portable Photosynthesis system, data represented by mean ± s.e.m, n = 7 for control and n = 8 for muscimol; statistical difference was determined by Student's *t-test*, **P< 0.05*.

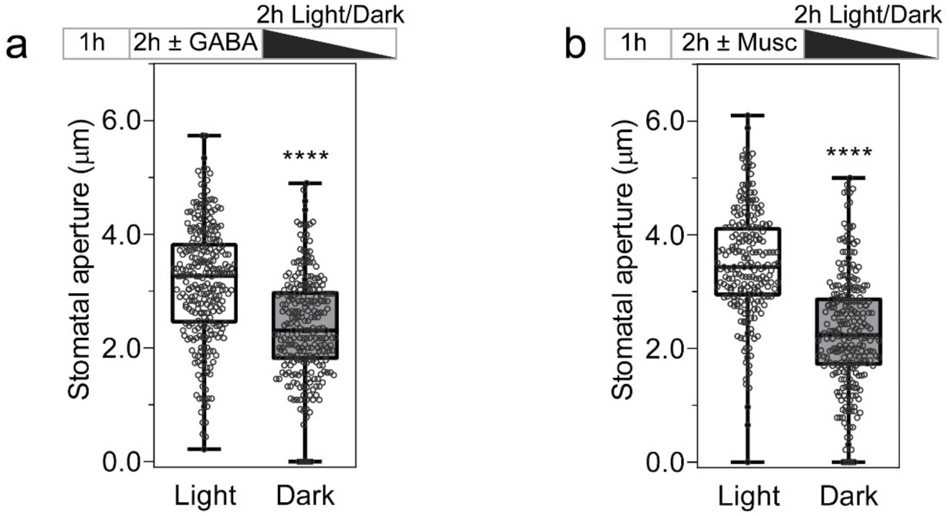


**Supplementary Figure 2. Guard cells are viable after treatment with GABA and muscimol. a-b,** Guard cells were competent in movement after removal of GABA or muscimol treatments, as open pores closed when exposed to dark following removal of GABA or muscimol. Epidermal strips were incubated under light for 1 h, followed by 2 h treatment of 2 mM GABA (**a**) or 10 μM muscimol (**b**), then epidermal strips were transferred into fresh stomatal measurement buffer with 2 h light or dark treatment before measurement; n = 275 for light and n = 248 for dark (**a**); n = 217 for light and n = 261 for dark (**b**); All data are plotted, statistical difference as determined by Students' *t*-test (**a, b**), **** P<*0.0001*.

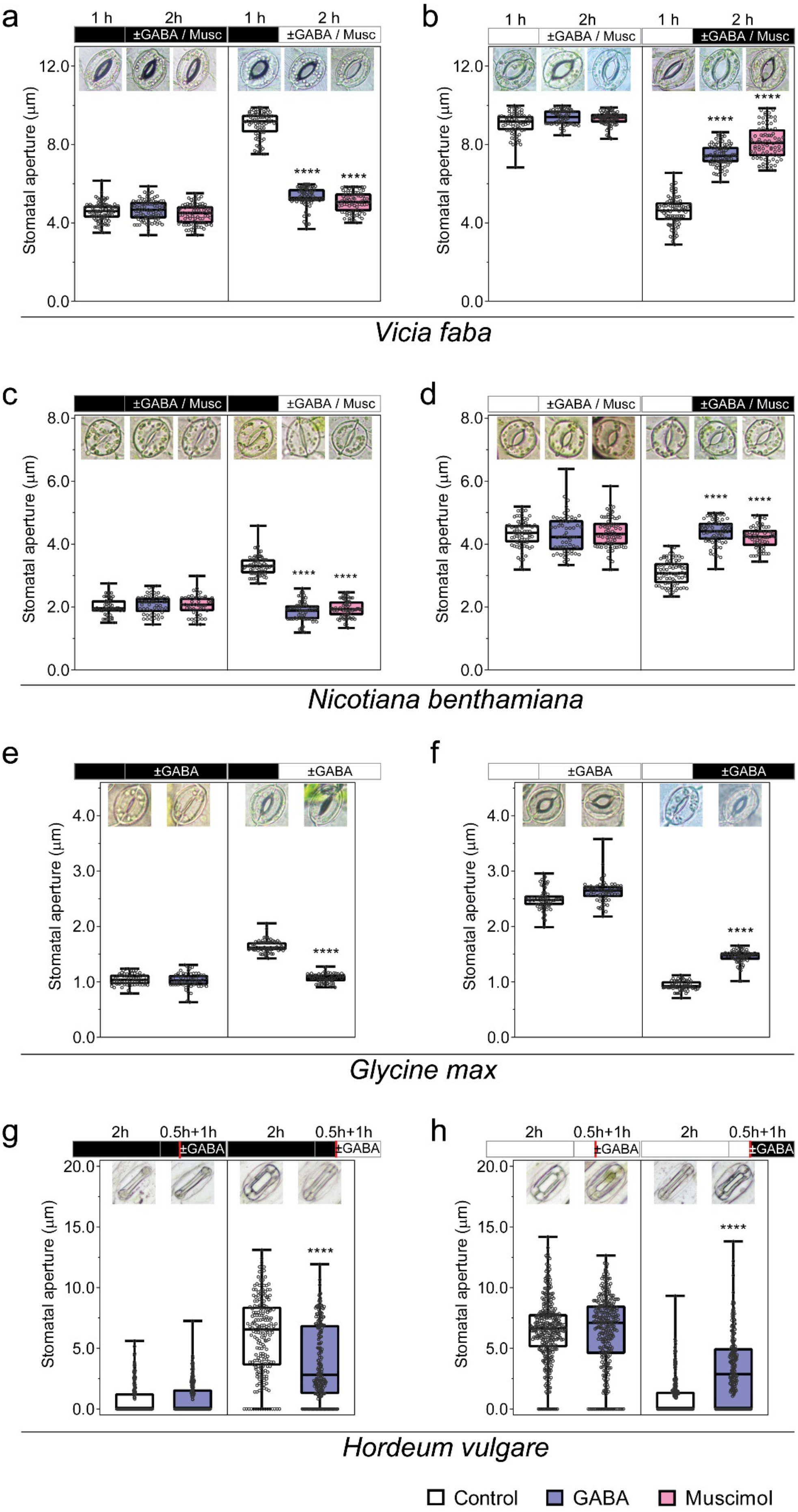
a
b
1 h
2h
2 h
±GABA / Musc
Stomatal aperture (µm)
12.0
8.0
4.0
0.0
****
Vicia faba
c
d
8.0
6.0
4.0
2.0
0.0
Nicotiana benthamiana
e
f
±GABA
4.0
3.0
2.0
1.0
0.0
Glycine max
g
h
0.5h+1h
20.0
15.0
10.0
5.0
0.0
Hordeum vulgare
Control
GABA
Muscimol

**Supplementary Figure 3**. **GABA and muscimol inhibit stomatal movement in response to light and dark in *Vicia faba* (broad bean), *Nicotiana benthamiana* (tobacco), *Glycine max* (soybean) and *Hordeum vulgare* (barley).** Epidermal strips were pre-incubated in stomatal measurement buffer for 1 h under dark (**a, c, e**) or light (**b, d, f**), followed by 2 h incubation under constant dark (**a, c, e**), light (**b, d, f**) as illustrated by black (dark) or white (light) bars, ± 2 mM GABA or 10 μM muscimol (Musc) as indicated; barley leaf samples were first detached and bathed in a modified measurement buffer under 2h dark (**g**) or light (100 μmol $m^{-2}$ $s^{-1}$) (**h**), then pre-treated in the fresh buffer ±1 mM GABA for 0.5 h as indicated by black or white bars; after this pre-treatment (break by red line), leaf samples were incubated in continuous dark (**g**), light (**h**), dark-to-light (**g**) or light-to-dark (**h**) transition for additional 1 h before the epidermal strips were peeled for imaging. n = 88 for control (constant dark), n = 85 for GABA (constant dark), n = 82 for muscimol (constant dark), n = 78 for control (dark-to-light transition), n = 106 for GABA (dark-to-light transition) and n = 76 for muscimol (dark-to-light transition) (**a**); n = 73 for control (constant light), n = 65 for GABA (constant light), n = 76 for muscimol (constant light), n = 89 for control (light-to-dark transition), n = 85 for GABA (light-to-dark transition) and n = 84 for muscimol (light-to-dark transition) (**b**); n = 50 for control (constant dark), n = 52 for GABA (constant dark), n = 50 for muscimol (constant dark), n = 63 for control (dark-to-light transition), n = 65 for GABA (dark-to-light transition) and n = 64 for muscimol (dark to light transition) (**c**); n = 73 for control (constant light), n = 60 for GABA (constant light), n = 78 for muscimol (constant light), n = 78 for control (light-to-dark transition), n = 67 for GABA (light-to-dark transition) and n = 65 for muscimol (light-to-dark transition) (**d**); n = 61 for control (constant dark), n = 60 for GABA (constant dark), n = 63 for control (dark-to-light transition) and n = 62 for GABA (dark-to-light transition) (**e**); n = 59 for control (constant light), n = 59 for GABA (constant light), n = 60 for control (light-to- dark transition) and n = 60 (dark-to-light transition) (**f**); n = 177 for control (constant dark), n = 220 for GABA (constant dark), n = 201 for control (dark-to-light transition) and n = 203 for GABA (dark-to-light transition) (**g**); n = 350 for control (constant light), n = 301 for GABA (constant light), n = 289 for control (light-to-dark transition) and n = 228 for GABA (light-to dark-transition) (**h**). All data are plotted points; statistical difference was determined by Two-way ANOVA, ****$P< 0.0001$.

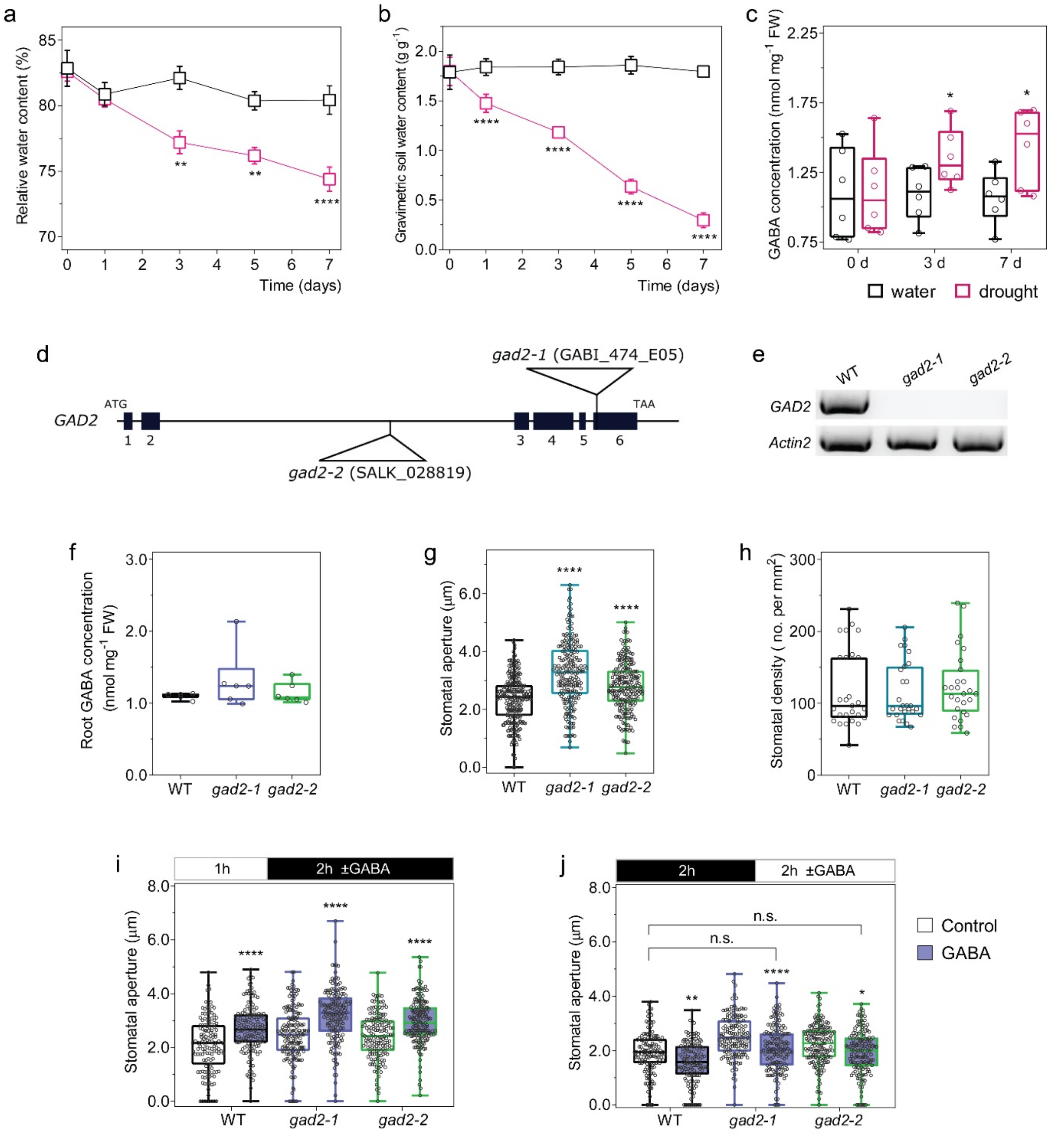


**Supplementary Figure 4. GABA accumulates in leaves of Arabidopsis upon drought, and *gad2* knockout plants have greater stomatal apertures but show wildtype (WT)-like responses to exogenous GABA and root GABA accumulation.** **a,** Relative water content in wildtype Arabidopsis leaves under well-water (black) or drought (red) treatments as indicated. **b,** Water content in the potted soil corresponding to the plants sampled in (**a**). **c,** Leaf GABA concentration of wildtype Arabidopsis following well-watered control treatment (black) or drought (red), sampled from (**a**); n = 6 plants (**a-c**). **d,** A diagram of *GAD2* T-DNA insertional mutant alleles in the Arabidopsis genome. **e,** Reverse transcriptional PCR semi-quantification of *GAD2* transcripts in Arabidopsis WT and *gad2* knockout plants, *Actin2* used as an internal control. **f,** Root GABA concentration of WT, *gad2-1* and *gad2-2* plants. Roots were harvested from 5-6 week-old plants grown hydroponically in basal nutrient solution (2 mM $NH_4NO_3$, 3 mM $KNO_3$, 0.1 mM $CaCl_2$, 2 mM KCl, 2 mM $Ca(NO_3)_2$, 2 mM $MgSO_4$, 0.6 mM $KH_2PO_4$, 1.5 mM NaCl, 50 μM NaFe(III)EDTA, 50 μM $H_3BO_3$, 5 μM $MnCl_2$, 10 μM $ZnSO_4$, 0.5 μM $CuSO_4$, 0.1 μM $Na_2MoO_3$, pH = 5.6 by KOH)[45], n = 6 plants. **g-j,** Stomatal aperture and density on the leaf abaxial side of Arabidopsis WT and *gad2* knockouts; epidermal strips were peeled and incubated in stomatal measurement buffer for 1 h under light before measurement n = 254 for WT, n = 215 for *gad2-1* and n = 226 for *gad2-2* (**g**); n = 27 sampling areas (0.57 x 0.42 mm) consisting of three areas per leaf, three leaves per plant and three plants per line sampled (**h**). **i-j,** Epidermal strips were pre-incubated in stomatal measurement buffer for 1 h under light (**i**) or 2 h dark (**j**), followed by 2 h incubation dark (**i**) or light (**j**) as indicated by black (dark) or white (light) bars ± blind treatment of 2 mM GABA or control; n = 135, 166 and 157 for WT, *gad2-1* and *gad2-2* with control treatment, n = 146, 162 and 174 for WT, *gad2-1* and *gad2-2* with GABA treatment (**i**); n = 139, 155 and 160 for WT, *gad2-1* and *gad2-2* with control treatment, n = 136, 155 and 153 for WT, *gad2-1* and *gad2-2* with GABA treatment (**j**). All data are plotted (**c, f-j**) or represented by mean ± s.e.m. (**a, b**); statistical difference was determined by Student's *t*-test (**c**), One-way ANOVA (**f-h**) or Two-way ANOVA (**a, b, i, j**), *$P$< 0.05, **$P$< 0.01 and ****$P$< 0.0001.

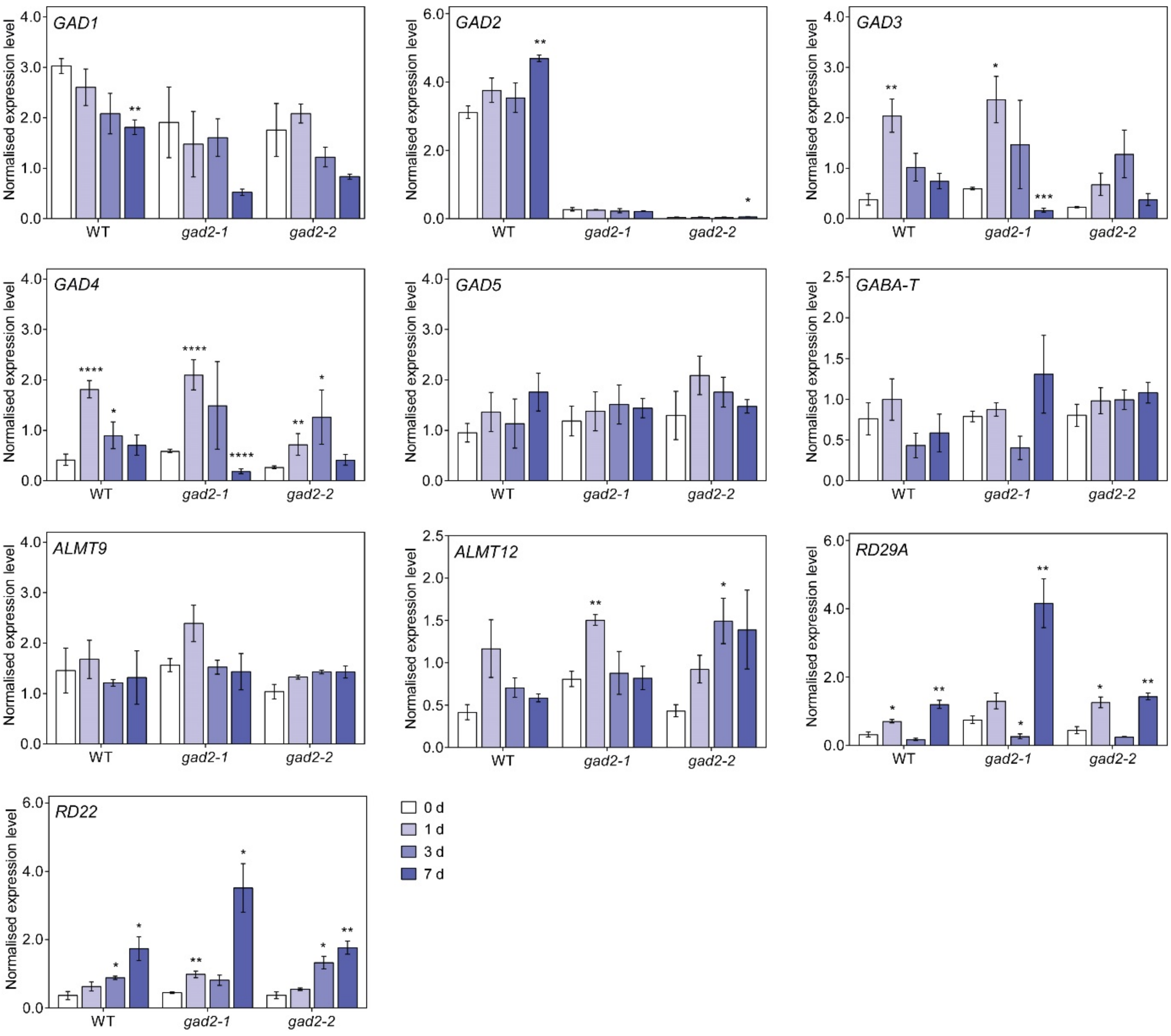


**Supplementary Figure 5. *gad2* knockouts have similar transcriptional profiles to wildtype plants of other *GADs*, *GABA-T*, *ALMT9*, *ALMT12* or ABA responsive genes under drought stress.** Quantitative real time PCR of *GAD1*, *GAD2*, *GAD3*, *GAD4*, *GAD5, GABA-T*, *ALMT9*, *ALMT12* and ABA marker genes –*RD29A* and *RD22*[46] in the leaves of Arabidopsis wildtype (WT), *gad2-1* and *gad2-2* plants following drought treatment for 0, 1, 3 and 7 days as indicated, expression levels were normalized against three control genes –*Actin2*, *EF1α* and *GAPDH-A*; data represented by means ± s.e.m; n = 3, statistical difference as determined via the comparison of genes from drought-treated plants (1, 3 and 7 days) with non-droughted (0 day) plants within the same genotype by Student's *t-test*, **P< 0.05* ***P< 0.01*, ****P< 0.001* and *****P< 0.0001*.

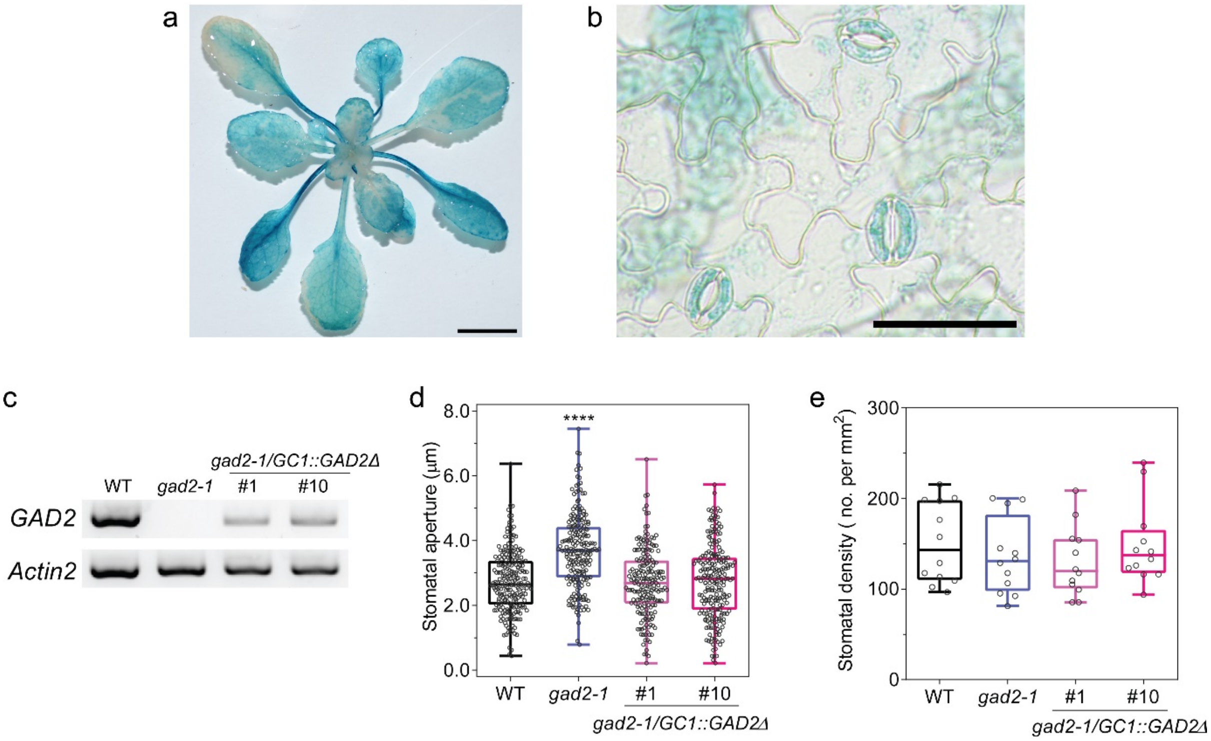


**Supplementary Figure 6. *GAD2* is highly expressed in leaves and guard cells, and guard-cell cell complementation of *GAD2Δ* restores stomatal aperture without modifying stomatal density. a-b,** GUS histochemical staining of *pGAD2::GUS* whole rosette**;** image of 3-4 week-old *pGAD2::GUS* plants after staining in histochemical staining buffer, scale bar = 5 mm (**a**) and epidermal peels from 3-4 week-old *pGAD2::GUS* leaves (**a**), scale bar = 50 μm (**b**). **c,** Reverse-transcriptional PCR quantification of *GAD2* transcripts in Arabidopsis wildtype (WT), *gad2-1*, *gad2-1/GC1::GAD2Δ* #1 and #10 plants. **d-e,** Stomatal aperture (**d**) and density (**e**) on the leaf abaxial side of Arabidopsis WT, *gad2-1*, *gad2-1/GC1::GAD2Δ* #1 and #10 plants; epidermal strips were peeled and incubated in stomatal measurement buffer for 1 h under light before measurement, n = 223 for WT, n = 212 for *gad2-1*, n = 197 for *gad2-1/GC1::GAD2Δ* #1 and n = 224 for *gad2-1/GC1::GAD2Δ* #10 (**d**); n = 12 leaf areas (0.57 x 0.42 mm); two areas per leaf, two leaves per plant and three plants per line were sampled (**e**). All data are plotted (**d, e**); statistical difference was determined by One-way ANOVA (**d, e**), ****$P$ <0.0001.

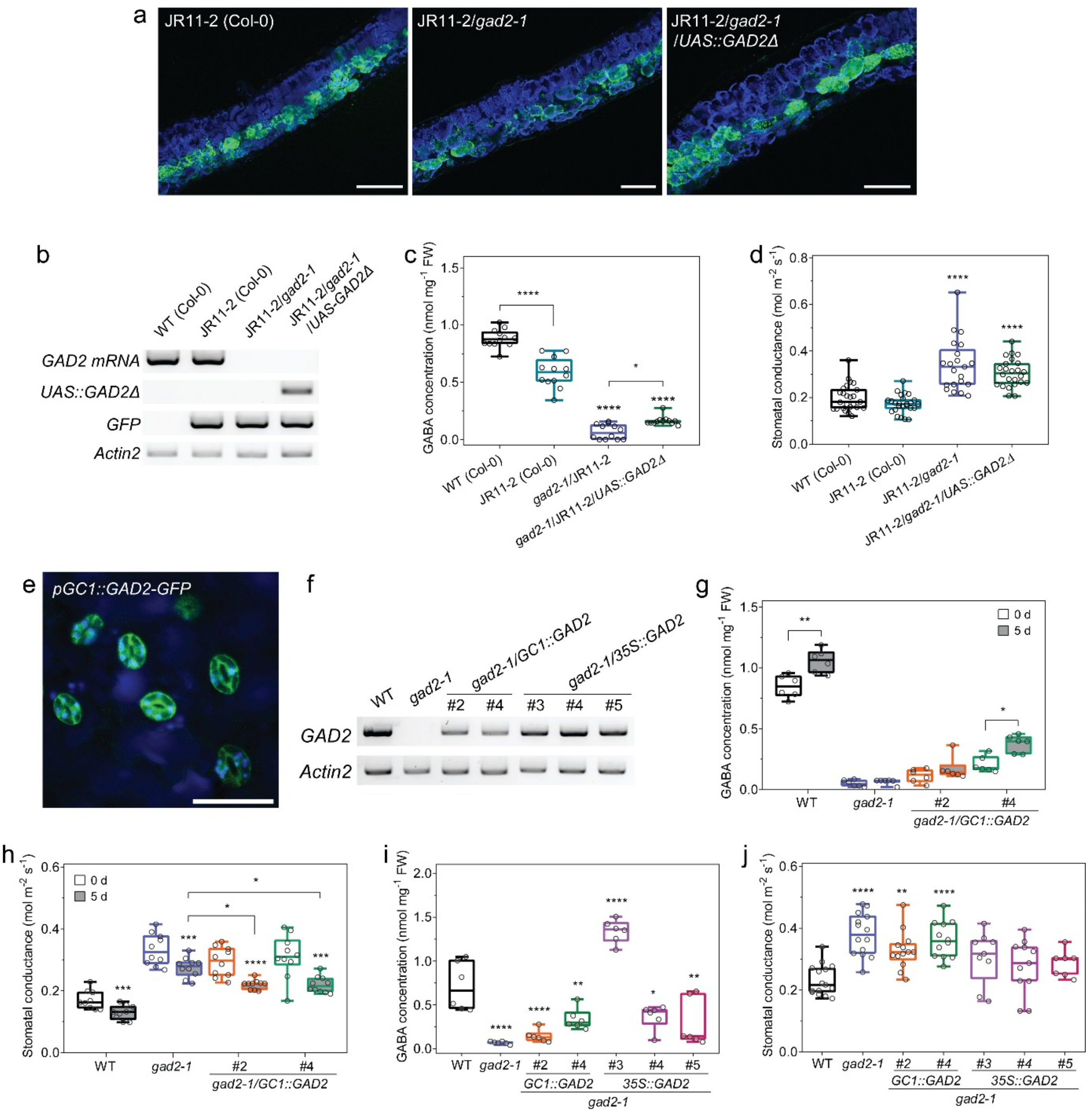


**Supplementary Figure 7. Spongy mesophyll-cell specific expression of *GAD2Δ* in *gad2-1* fails to restore stomatal conductance back to wildtype (WT) levels, and guard-cell specific expression of full-length *GAD2* only reduces the stomatal conductance of *gad2* knockout plants under drought. a,** Images of leaf transverse cross-sections (30 μm thickness) of 3-4 week-old segregated mesophyll-specific enhancer-trap line JR11-2[29] backcrossed into Arabidopsis Col-0 background[38], JR11-2 in *gad2-1* background (JR11-2/*gad2-1*) and JR11-2/*gad2-1* expressing *UAS::GAD2Δ* (JR11-2/*gad2-1*/*UAS::GAD2Δ*), scale bars = 100 μm. GFP fluorescence shown in green indicates cells in which *GAD2* expression will be activated by the yeast transcription factor GAL4, blue indicates chlorophyll autofluorescence. **b,** Reverse-transcriptional PCR quantification of native *GAD2* mRNA (*GAD2mRNA*), *GAD2Δ* driven by *UAS* element (*UAS::GAD2Δ*), *GFP* and *Actin2* transcripts in Arabidopsis WT (Col-0), JR11-2 (Col-0), JR11-2/*gad2-1* and JR11-2/*gad2-1*/*UAS::GAD2Δ* plants, *Actin2* used as an internal control. **c-d,** Leaf GABA concentration (**c**) and stomatal conductance (**d**) of 5-6 week-old Arabidopsis WT (Col-0), JR11-2 (Col-0), JR11-2/*gad2-1* and JR11-2/*gad2-1*/*UAS::GAD2Δ* plants, stomatal conductance was measured by AP4 Leaf Porometer (**d**); n = 12 (**c**); n = 25 for WT and JR11-2, n = 21 for JR11-2/*gad2-1* and n = 24 for JR11-2/*gad2-1*/*UAS::GAD2Δ* (**d**). **e,** Confocal image of *gad2*-1 leaves expressing full-length *GAD2* tagged with *GFP* driven by *GC1* promoter (*GC1::GAD2-GFP*), scale bar = 50 μm. **f,** Reverse-transcriptional PCR quantification of *GAD2* transcripts in wildtype (WT), *gad2-1* and *gad2-1* complementation with full-length *GAD2* driven by guard-cell promoter *GC1* (*gad2-1/GC1::GAD2* #2 and #4) or by a pro35S-CAMV constitutive promoter (*gad2-1/35S::GAD2* #3, #4 and #5), *Actin2* used as an internal control. **g-h,** Leaf GABA concentration and stomatal conductance of WT, *gad2-1*, *gad2-1/GC1::GAD2* #2 and #4 plants; n = 6 plants for GABA measurement before (0 d) and after drought treatment for 5 days (5 d) as indicated (**g**); The stomatal conductance of 5-6 week-old plants was determined by AP4 Leaf Porometer 0 d and 5 d after drought treatment, n = 9 for WT and n = 10 for *gad2-1*, *gad2-1/GC1::GAD2* #2 and #4 (**h**). **i-j,** Leaf GABA concentration and stomatal conductance of WT, *gad2-1*, *gad2-1/GC1::GAD2* #2, #4, *gad2-1/35S::GAD2* #3, #4 and #5 plants; n = 6 plants (**i**); the stomatal conductance of WT, *gad2-1*, and *gad2-1* complementation lines; stomatal conductance of 5-6 week-old plants was determined by AP4 Leaf Porometer in normal conditions, n = 15 for WT, n = 14 for *gad2-1*, n = 13 for *gad2-1/GC1::GAD2* #2, n = 12 for *gad2-1/GC1::GAD2* #4, n = 9 for *gad2-1/35S::GAD2* #3, n = 11 for *gad2-1/35S::GAD2* #4 and n = 7 for *gad2-*

*1/35S::GAD2* #5 (**j**). All data are plotted (**c, d, g-j**); statistically differences were determined by One-way ANOVA by comparing with either JR11-2 (**c, d**) or WT (**i, j**), Student's t-test (**g**) within genotypes or by Two-way ANOVA (**h**), **P*< 0.05, ***P*< 0.01, ****P* < 0.001 and *****P*< 0.0001.

a

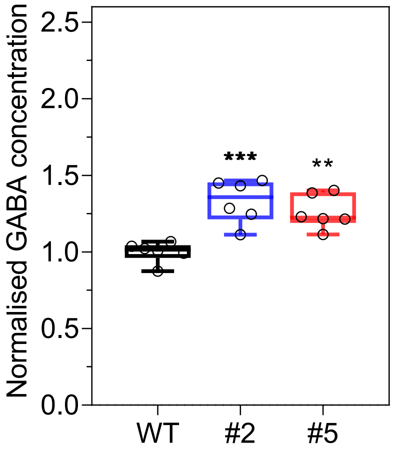


b

WT
(22.2%)

*GC1::GAD2Δ* #2
(41.7%)

*GC1::GAD2Δ* #5
(78.6%)

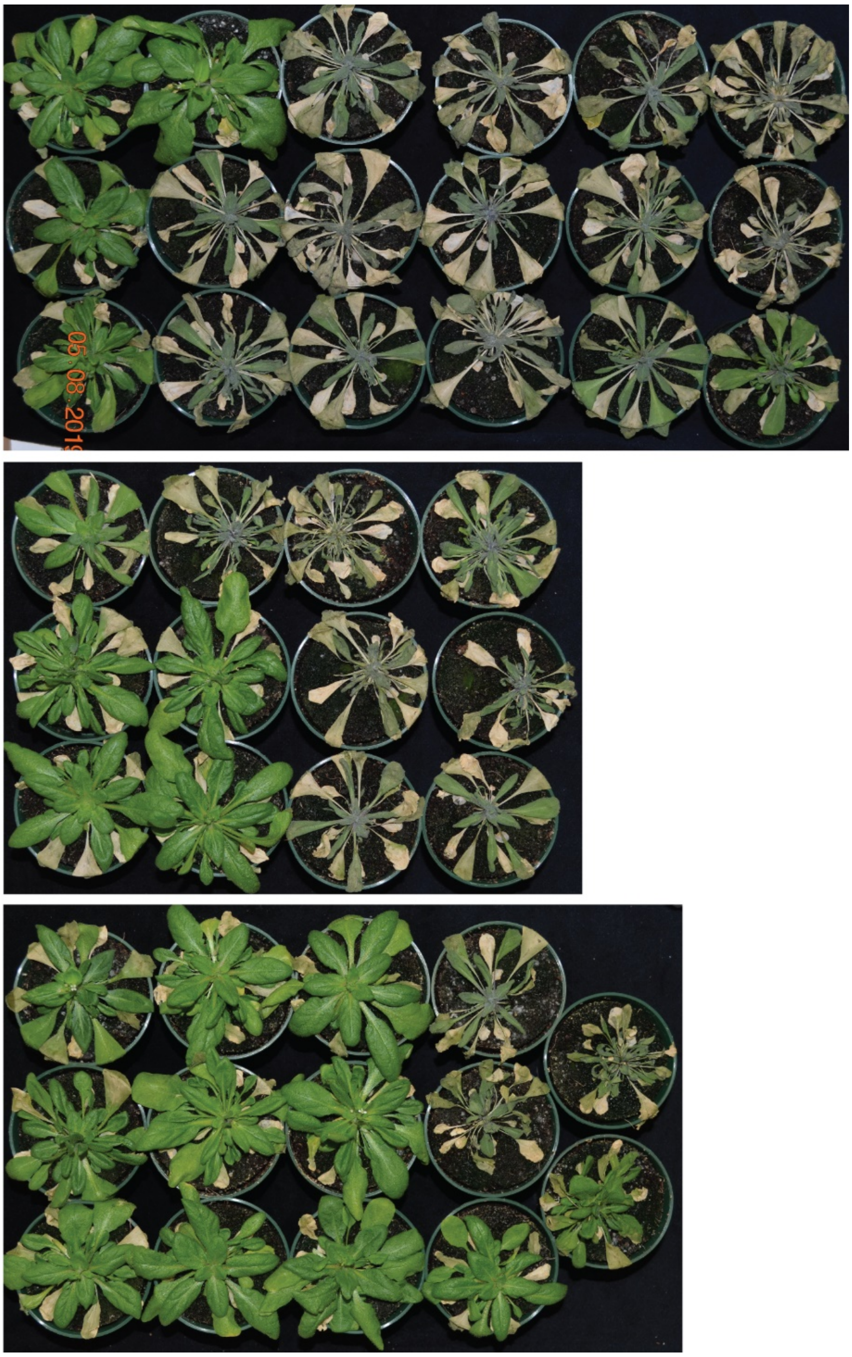

c

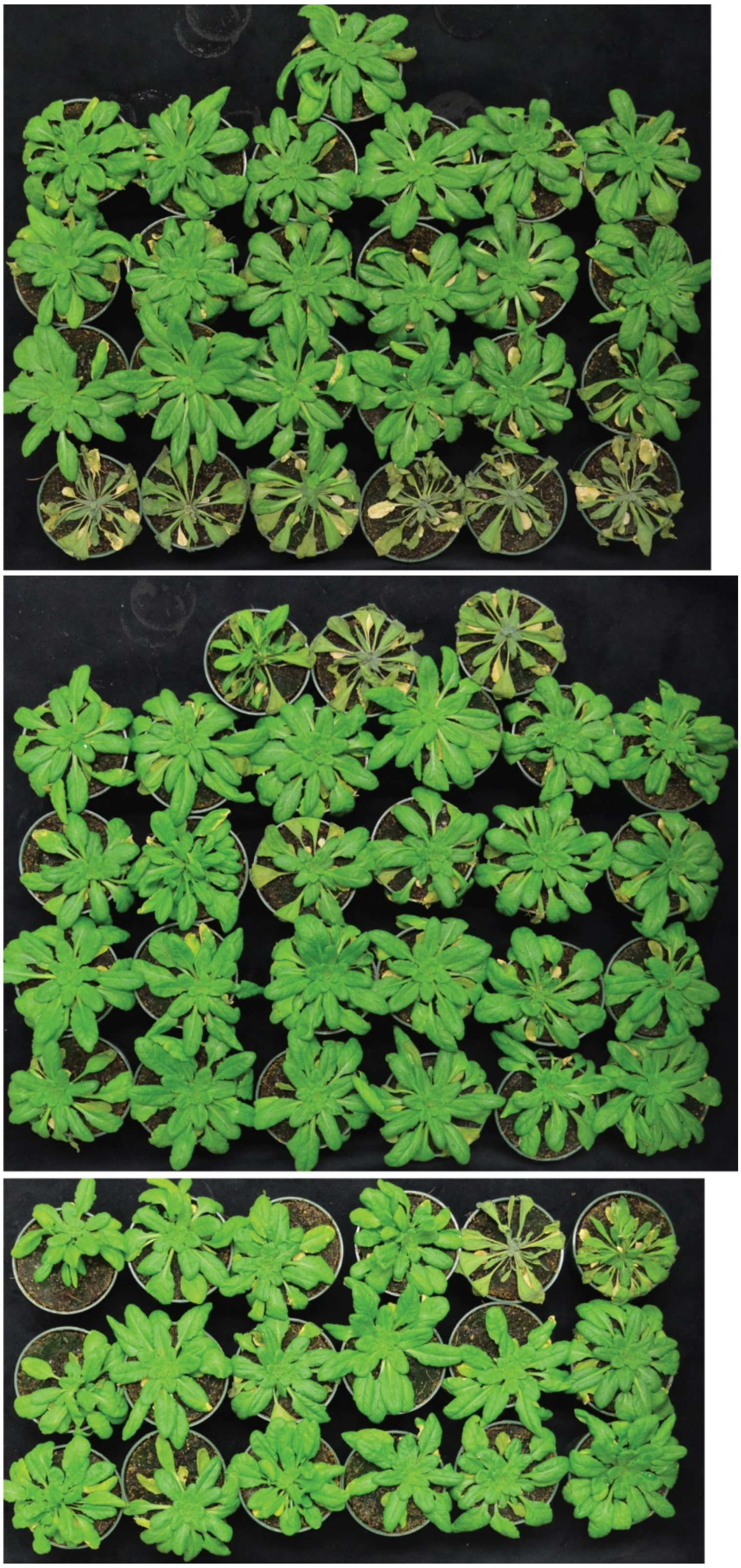

WT
(77.8%)

*GC1::GAD2Δ* #2
(88.9%)

*GC1::GAD2Δ* #5
(88.9%)

d

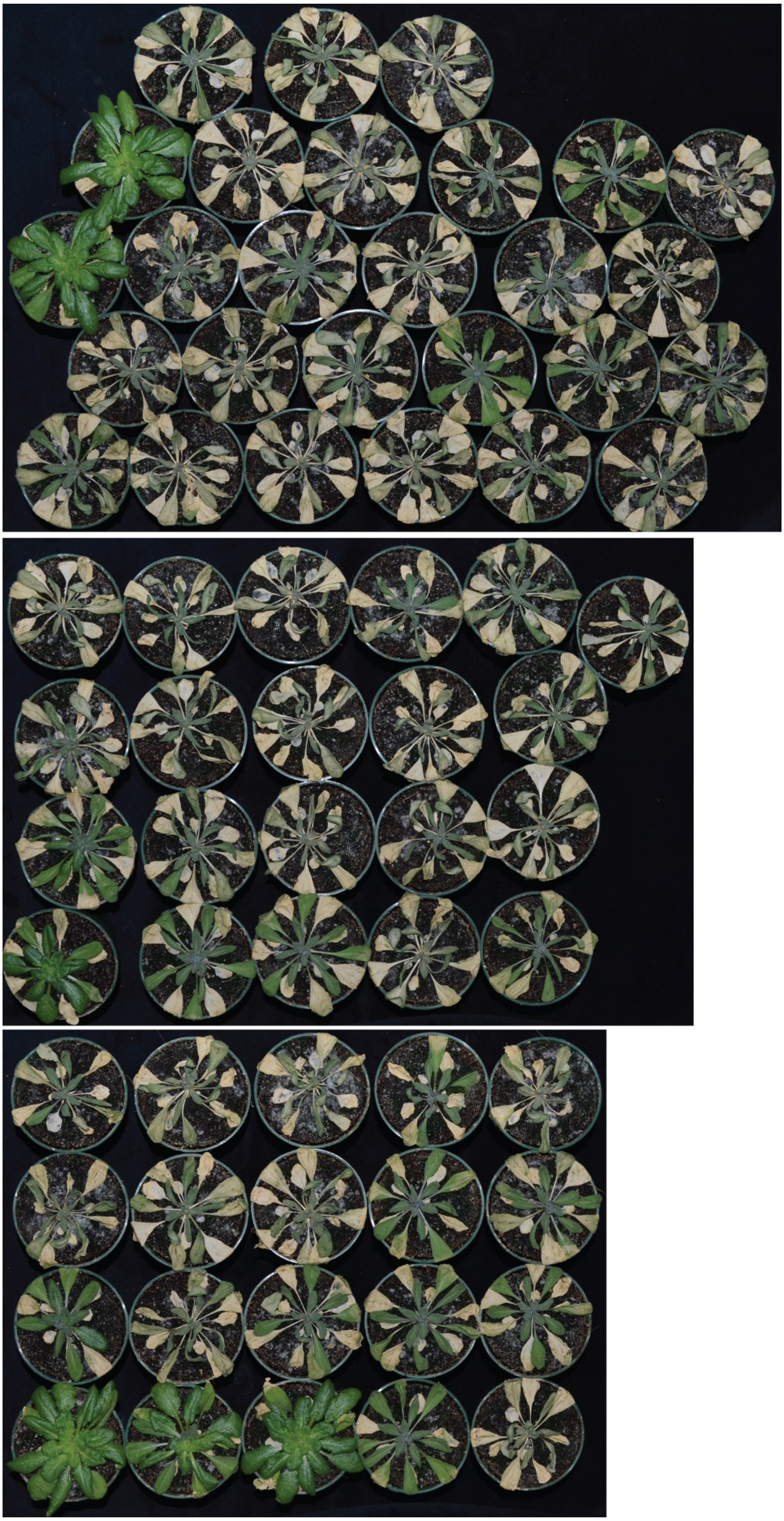

WT
(11.1%)

*GC1::GAD2Δ* #2
(19.0%)

*GC1::GAD2Δ* #5
(35%)

**Supplementary Figure 8. Leaf GABA accumulation and recovery of re-watered wildtype, plants *GC1::GAD2Δ* #2 and #5 from drought treatment. a**, Normalised GABA accumulation in the leaves of 5-6 week-old Arabidopsis wildtype, *GC1::GAD2Δ* #2 and #5 plants; n = 6, statistical analysis was determined by One-way ANOVA, **$P<0.01$ and **$P<0.001$. **b-d**, Re-water recovery of wildtype, *GC1::GAD2Δ* #2 and #5 plants from drought in three different batches of experiments, plants were re-watered at 2 days after all plant wilting by drought. A higher recovery rate of *GC1::GAD2Δ* #2 and #5 plants than WT was observed from re-watering in all three experiments (**b-d**); 4 out of 18 (22%) wildtype, 5 out of 12 (41.7%) *GC1::GAD2Δ* #2 and 11 out of 14 (78.6%) *GC1::GAD2Δ* #5 plants were recovered from re-water in the first trail (**b**); 21 out of 27 (77.8%) wildtype, 24 out of 27 (88.9%) *GC1::GAD2Δ* #2 and 16 out of 18 (88.9%) *GC1::GAD2Δ* #5 plants were recovered from re-water in the second trail (**c**); 3 out of 27 (11.1%) wildtype, 4 out of 21 (19.0%) *GC1::GAD2Δ* #2 and 7 out of 20 (35%) *GC1::GAD2Δ* #5 plants were recovered from re-water in the third trail (**d**).

.

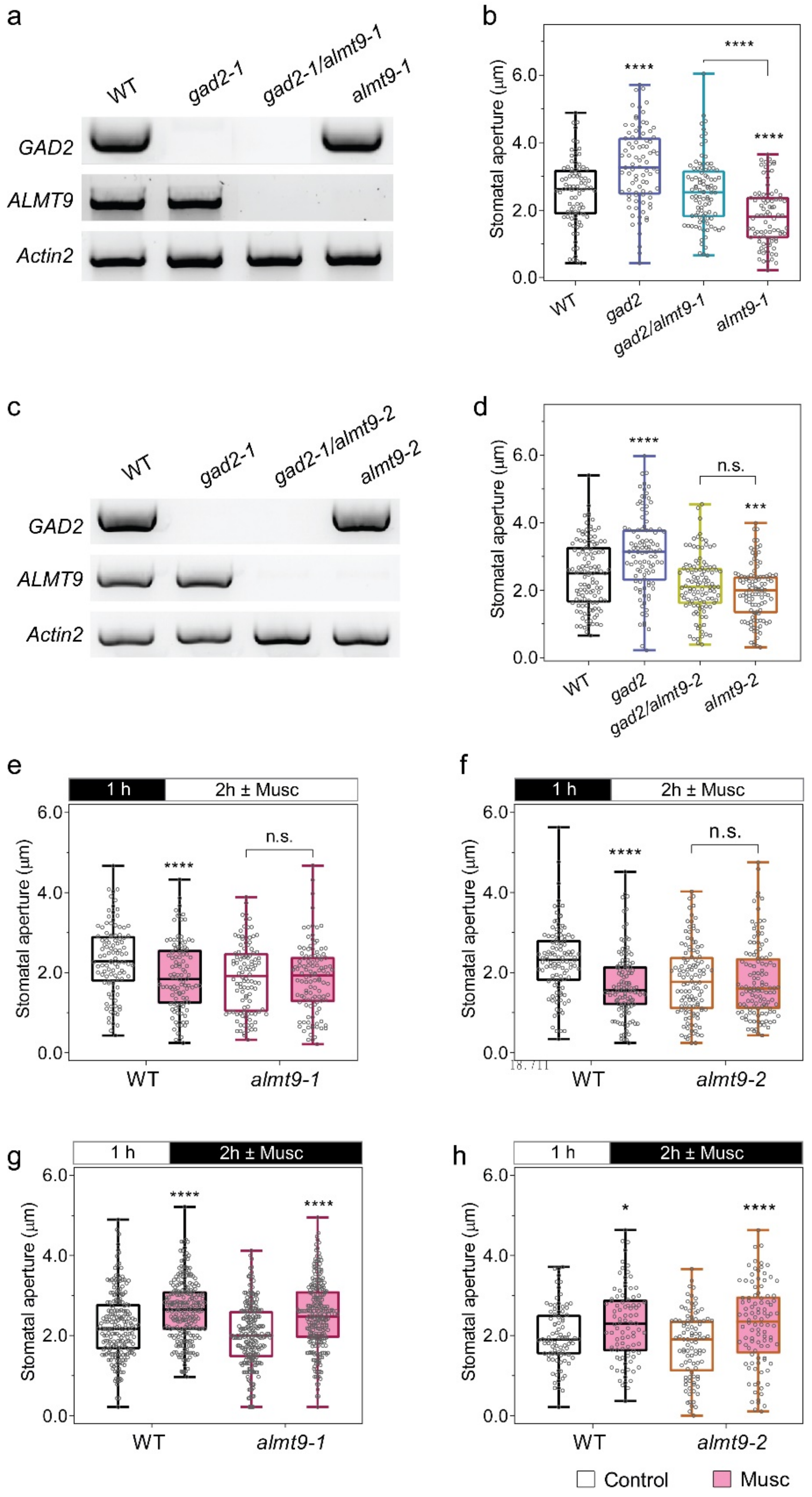

a
WT
gad2-1
gad2-1/almt9-1
almt9-1
GAD2
ALMT9
Actin2
b
Stomatal aperture (μm)
WT
gad2
gad2/almt9-1
almt9-1
c
WT
gad2-1
gad2-1/almt9-2
almt9-2
GAD2
ALMT9
Actin2
d
Stomatal aperture (μm)
n.s.
WT
gad2
gad2/almt9-2
almt9-2
e
1 h
2h ± Musc
Stomatal aperture (μm)
n.s.
WT
almt9-1
f
1 h
2h ± Musc
Stomatal aperture (μm)
n.s.
WT
almt9-2
g
1 h
2h ± Musc
Stomatal aperture (μm)
WT
almt9-1
h
1 h
2h ± Musc
Stomatal aperture (μm)
WT
almt9-2
Control
Musc


18.711

**Supplementary Figure 9**. ***almt9*** **knockouts abolish muscimol-inhibition of stomatal opening but does not affect closure, and *almt9/gad2* rescues the larger stomatal aperture of *gad2* knockout plants.** **a,** Reverse-transcriptional PCR quantification of *GAD2*, *ALMT9* and *Actin2* transcripts in Arabidopsis wildtype (WT), *gad2-1*, *gad2-1/almt9-1* and *almt9-1* plants, *Actin2* used as an internal control. **b,** Stomatal aperture of WT (n = 86), *gad2-1* (n = 86), *gad2-1/GC1::GAD2Δ #10* (n = 86), *gad2-1/almt9-1* (n = 95) and *almt9-1* (n = 87) plants, epidermal strips were peeled and incubated in stomatal measurement buffer for 2 h under light before measurement. **c,** Reverse-transcriptional PCR quantification of *GAD2*, *ALMT9* and *Actin2* transcripts in WT, *gad2-1*, *gad2-1/almt9-2* and *almt9-2* plants, *Actin2* used as an internal control. **d,** Stomatal aperture of WT (n = 115), *gad2-1* (n = 100), *gad2-1/almt9-2* (n = 106) and *almt9-2* (n = 104) plants; epidermal strips were incubated under light for 2 h before measurement. **e-h**, Stomatal aperture of wildtype (WT) and *almt9* knockout plants in response to dark or light. Epidermal strips were pre-incubated in stomatal measurement buffer for 1 h under dark (**e, f**) or light (**g, h**), followed by light (**e, f**) or dark (**g, h**) for 2 h as indicated by black (dark) or white (light) bars above graphs, ± 10 µM muscimol (Musc); n = 105 for WT and n = 106 for *almt9-1* with control treatment, n = 106 for WT and n = 111 for *almt9-1* with muscimol treatment (**e**); n = 88 for wildtype (WT) (control); n = 108 for WT (control), n = 116 for *almt9-2* (control), n = 119 for WT (muscimol) and n = 121 for *almt9-2* (muscimol) (**f**); n = 210 for WT and n = 233 for *almt9-1* with control treatment, n = 240 for WT and n = 245 for *almt9-1* with muscimol treatment (**g**); n = 88 for WT (control), n = 96 for *almt9-2* (control), n = 90 for WT (muscimol) and n = 100 for *almt9-2* (muscimol) (**h**); all experiments were repeated at least twice from different batches of plants with blind treatments (**e-h**)..

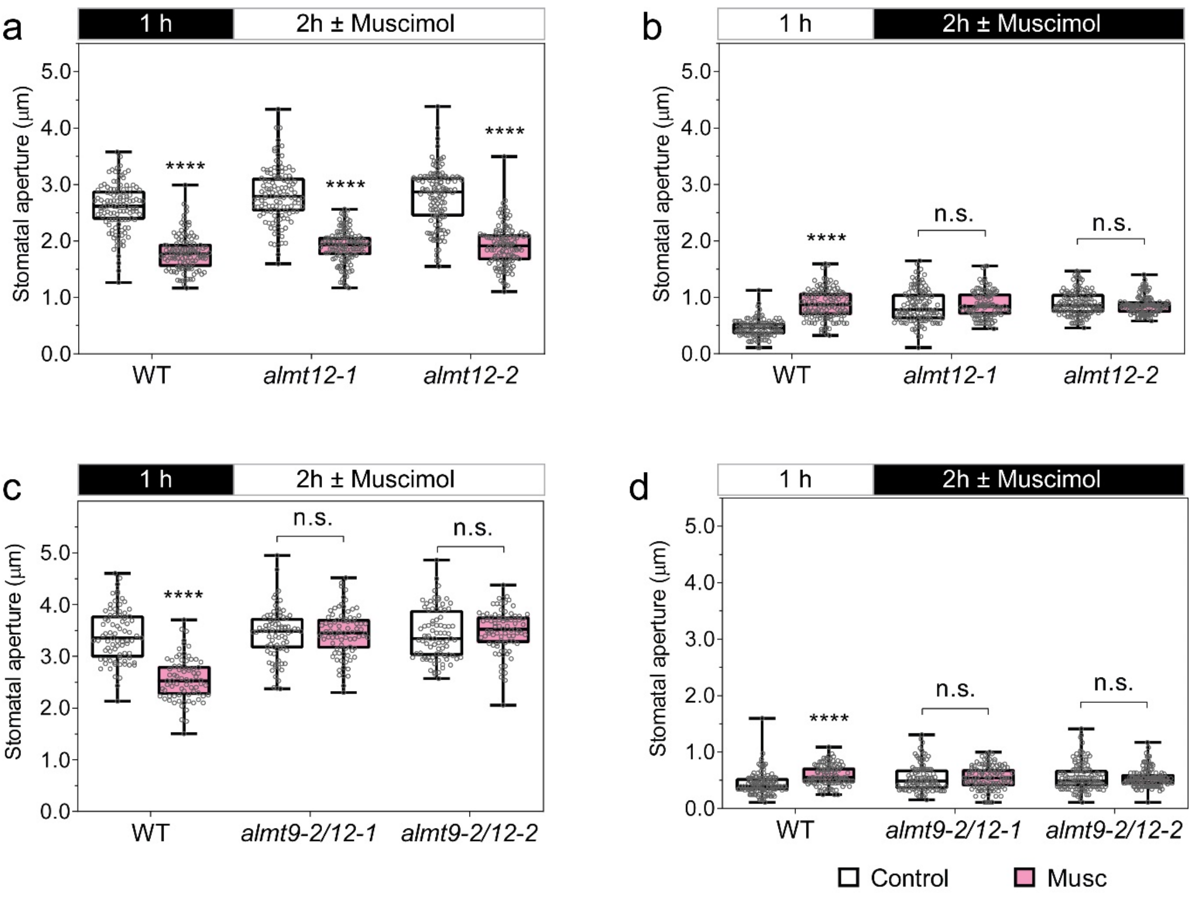


**Supplementary Figure. 10. Stomatal aperture measurement of wildtype (WT), *almt12* and *almt9-2/almt12* knockout plants in response to dark or light.** Epidermal strips were pre-incubated in stomatal measurement buffer for 1 h in the dark (**a, c**) or light (**b, d**), followed by 2 h incubation in the light (**a, c**) or dark (**b, d**) as indicated by black (dark) or white (light) bars above the plots ± 2 mM GABA or 10 µM muscimol (Musc); n = 116 for WT (control), n = 119 for *almt12-1* (control), n = 120 for *almt12-2* (control), n = 117 for WT (Musc), n = 122 for *almt12-1* (Musc) and n = 116 for *almt12-2* (Musc) (**a**); n = 105 for WT (control), n = 115 for *almt12-1* (control), n = 122 for *almt12-2* (control), n = 122 for WT (Musc), n = 107 for *almt12-1* (Musc) and n = 118 for *almt12-2* (Musc) (**b**); n = 78 for WT (control), n = 82 for *almt9-2/12-1* (control), n = 83 for *almt9-2/12-2* (control), n = 75 for WT (Musc), n = 81 for *almt9-2/12-1* (Musc) and n = 81 for *almt9-2/12-2* (Musc) (**c**); n = 114 for WT (control), n = 104 for *almt9-2/12-1* (control), n = 120 for *almt9-2/12-2* (control), n = 107 for WT (Musc), n = 106 for *almt9-2/12-1* (Musc) and n = 127 for *almt9-2/12-2* (Musc) (**d**). All data are plotted, statistical difference was determined using Two-way ANOVA, ****P* < 0.001, *****P* < 0.0001; all experiments were repeated at least twice from different batches of plants with blind treatments (**a-d**).
